# Dynamic Scan Procedure for Detecting Rare-Variant Association Regions in Whole Genome Sequencing Studies

**DOI:** 10.1101/552950

**Authors:** Zilin Li, Xihao Li, Yaowu Liu, Jincheng Shen, Han Chen, Hufeng Zhou, Alanna C. Morrison, Eric Boerwinkle, Xihong Lin

## Abstract

Whole genome sequencing (WGS) studies are being widely conducted to identify rare variants associated with human diseases and disease-related traits. Classical single-marker association analyses for rare variants have limited power, and variant-set based analyses are commonly used to analyze rare variants. However, existing variant-set based approaches need to pre-specify genetic regions for analysis, and hence are not directly applicable to WGS data due to the large number of intergenic and intron regions that consist of a massive number of non-coding variants. The commonly used sliding window method requires pre-specifying fixed window sizes, which are often unknown as *a priori*, are difficult to specify in practice and are subject to limitations given genetic association region sizes are likely to vary across the genome and phenotypes. We propose a computationally-efficient and dynamic scan statistic method (Scan the Genome (SCANG)) for analyzing WGS data that flexibly detects the sizes and the locations of rare-variants association regions without the need of specifying a prior fixed window size. The proposed method controls the genome-wise type I error rate and accounts for the linkage disequilibrium among genetic variants. It allows the detected rare variants association region sizes to vary across the genome. Through extensive simulated studies that consider a wide variety of scenarios, we show that SCANG substantially outperforms several alternative rare-variant association detection methods while controlling for the genome-wise type I error rates. We illustrate SCANG by analyzing the WGS lipids data from the Atherosclerosis Risk in Communities (ARIC) study.

## Introduction

An important goal of human genetic research is to identify the genetic basis for human diseases and traits. Genome-wide association studies (GWAS) have been widely used to dissect the genetic architecture of complex diseases and quantitative traits in the past ten years. Although GWAS has been successful for identifying thousands of common genetic variants putatively harboring susceptibility alleles for complex diseases,^1^ these common variants only explain a small fraction of heritability^2^ and the vast majority of variants in the human genome are rare.^3; 4^ In recent years, studies based on Whole Exome Sequencing have been conducted to detect rare variant associations in the coding sequences.^5; 6^ However, the majority of rare variants are non-coding and are located in introns and intergenic regions that are not covered by whole exome sequencing. The ENCODE project showed that a large fraction of these regions are functionally active,^7^ suggesting that these regions may also be linked with diseases or traits. A rapidly increasing number of Whole Genome Sequencing (WGS) association studies are being conducted to identify rare variants associated with complex traits and diseases. Examples include the Genome Sequencing Program of the National Human Genome Research Institute (NHGRI), and the Trans-Omics for Precision Medicine (TOPMed) Program of the National Heart, Lung, and Blood Institute (NHLBI).

The common analytic strategy in GWAS is to perform individual variant tests followed by a multiple testing adjustment to identify trait- or disease-associated variants. However, individual variant analysis is not applicable for analyzing rare variants due to a lack of power.^8–10^ There is an active recent literature on rare variant analysis methods which jointly test the effects of multiple variants in a pre-specified genetic set, such as burden tests,^11–13^ and non-burden tests^14–17^ such as Sequence Kernel Association Test (SKAT^14^). Instead of testing each variant individually, these tests evaluate the cumulative effects of multiple variants in a gene, and can boost power when multiple variants in the region are associated with a disease or a trait.^18; 19^ A limitation of these variant set-based tests is that they need to pre-specify genetic regions to be used for analysis. Hence these existing gene-based approaches are not directly applicable to analysis of all the variants in WGS data, as variant sets are not well defined across the genome.

Sliding window methods have been proposed to identify regions of the genome harboring rare variants associated with complex traits and diseases. These procedures perform WGS association analysis by pre-specifying a certain size of moving windows that begin at the beginning of each chromosome with a fixed skip length and move along the genome.^20–22^ A major limitation of the sliding window approach is that it requires pre-specifying a fixed window size, either using the base pairs or number of variants, which is often unknown in advance and might not be optimal across the genome and phenotypes. The sliding window size is often difficult to specify in practice, since the exact sizes of trait or disease-associated regions might differ across the genome and vary with phenotypes. Indeed, the sliding window methods are likely to lose power if a pre-specified region is too big by including too many neutral variants, e.g., by sub-regions that contain only neutral variants, or a pre-specified region is too small by excluding adjacent regions that contain association signals. It is hence of substantial interest to develop a dynamic procedure that can scan the genome continuously by allowing the sizes and locations of the units for rare variant association testing to vary across the genome so that one can flexibly identify the associated regions without specifying a fixed window size and location as *a priori*.

We propose using scan statistic-based methods to scan the whole genome continuously by allowing for overlapping windows of different sizes by “shifting forward” a window of a given size with a small number of variants at a time and searching for the windows containing association signals. Scan statistics are a broad class of methods looking for clusters of events in time and space. Some of them have been successfully used in genetic studies, e.g., to refine the search for new genes in linkage studies^23^ and to identify chromosomal patterns of SNP association for common variants in GWAS.^24; 25^ Recently, likelihood-ratio based scan statistic procedures^26; 27^ have been proposed for identifying clusters of rare-disease variants in a large genetic region, such as a gene. However, these likelihood-ratio based scan methods are designed for refining a disease clustering region in a gene, but not for testing associations across the genome. Further, current methods do not allow covariates (e.g. age, sex, and population structures), and can only be used for binary traits. There is a pressing need to develop powerful scan methods for testing rare variant associations in WGS studies that can dynamically detect the varying sizes of rare variant association regions and their locations across the genome.

In this paper, we develop the SCAN the Genome (SCANG) method, which is a flexible and computationally efficient scan statistic procedure that uses the p-value of a variant set-based test as a scan statistic of each moving window, to detect rare variant association regions for both continuous and dichotomous traits. The goal of SCANG is to detect whether any rare-variant association region exists across the genome, and if they do exist, to identify the locations and sizes of these association regions. Specifically, SCANG first fits the null linear or logistic model that includes covariates, e.g., age, sex and ancestry PCs, but no genetic variants. Second, SCANG applies set-based tests to all possible candidate moving windows of different sizes within a pre-specified window range of practical interest. We include three tests in the SCANG framework: the burden test (SCANG-B), SKAT (SCANG-S) and an efficient omnibus test to aggregate information of the burden test and SKAT and different choices of weights using the ACAT method ^28; 29^ (SCANG-O). All the set-based tests share the same reduced model under the null hypothesis and hence the fitted null model is the same and only needs to be fit once when scanning the genome. Therefore, the computation of SCANG is highly efficient. Third, SCANG generates an empirical threshold calculated by Monte Carlo simulation, to control the Genome-wise/Family-wise Type I Error Rate (GWER/FWER) at a given level, e.g., 0.05. The windows with the p-values smaller than this threshold are detected as genome-wise significant association regions. Both individual-window p-values and the genome-wise/family-wise p-values of these genome-wise significant windows are given.

We demonstrate through simulation that SCANG is often more powerful than existing methods across a broad range of study designs for both continuous and dichotomous traits. We also apply SCANG to the analysis of the WGS and lipid traits from the Atherosclerosis Risk in Communities (ARIC) study. By allowing for estimating the optimal sizes of the genome-wide significant variants-phenotype association regions, SCANG detected a significant association between rare variants in a 4,637 bp region which resides in *NECTIN2* [MIM:600798] on chromosome 19 with small dense low-density lipoprotein cholesterol (sdLDL-c), an association that is missed by the conventional sliding window approach.^20^ SCANG also detected more association regions between rare variants and lipoprotein(a) (Lp(a)) than the sliding window procedure for all the considered tests.^20^

## Material and Methods

### SCAN the Genome (SCANG)

SCANG is a dynamic adaptive scan procedure for detecting regions of rare variants that are significantly associated with a phenotype. For each region, SCANG analytically calculates a set-based p-value and also empirically calculates a genome-wise p-value, which adjusts for multiple testing of all the moving windows of different sizes under consideration, including some overlapping windows, across the genome.

#### Aggregation Tests for Multiple Variants of a Given Region

Suppose that the data are from *n* subjects and there are *p* variants across the genome. Given a genetic set with *p*_0_ variants, for subject *i*, let *y_i_* denote a phenotype with mean *μ_i_*, ***X_i_*** = (*X*_*i*1_, *X*_*i*2_,…,*X_iq_*) denote the covariates, and ***G_i_*** = (*G*_*i*1_…, *G_im_*) be a vector of *m* variants in a given variant set. To relate the sequenced variants in the set to the phenotype, when samples are unrelated, we consider the following Generalized Linear Model (GLM):

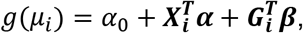

where *g*(*μ*) = *μ* for a continuous trait, and *g*(*μ*) = logit(*μ*) for a binary trait, *α*_0_ is an intercept, ***α*** = (*α*_1_,…, *α_q_*)^T^ is a *q* x 1 column vector of covariate effects and ***β*** = (*β*_1_,…, *β_m_*)^T^ is a *m* x 1 vector of the genotype effects in the variant set. Under GLMs, the null hypothesis of no association between the genetic variants in a set with the trait, adjusting for covariates and population structure and relatedness, corresponds to testing the null hypothesis *H*_0_: ***β = 0***, that is, *β*_1_ = *β*_2_ = ⋯ = *β*_*p*_0__ = 0. The score statistic of the marginal model for variant *j* is defined as

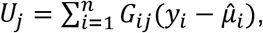

where 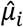 is the estimated mean of *y_i_* under the global null hypothesis (*H*_0_: ***β*** = 0) and is obtained by fitting the global null model *g*(*μ_i_*) = *α*_0_ + ***X_i_α***. For an extension to account for relatedness using Generalized Linear Mixed Models (GLMMs), see the Discussion Section.

To jointly test the effects of multiple variants in a given region *I*, burden tests collapse genotype information of all the variants in the region into a single genotype score and test for the association between this score and a trait. Specifically, burden tests count the total number of minor alleles in the set and the corresponding score statistic is

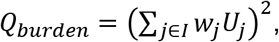

where *w_j_* is the weight for variant *j*.^10^ *Q_burden_* asymptotically follows a chi-square distribution with 1 degree of freedom and its p-value can be computed analytically.

Another widely used set-based test is SKAT,^14^ which tests for the genetic set association using a variance-component score test. The corresponding SKAT test statistic is

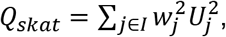

where *wj* is the weight for variant *j*.^14^ *Q_skat_* asymptotically follows a mixture of chi-square distributions and its p-value can also be obtained analytically.^14^

For both burden test and SKAT, we consider using the family of beta densities of MAF as weights,^14^ i.e., for each variant *j, w_j_* = *Beta*(*MAF_j_; a*_1_,*a*_2_), the beta density function with two parameters *a*_1_ and *a*_2_. Common choices of the parameters are *a*_1_ = 1 and *a*_2_ = 1, which correspond to equal weights under the assumption of the same effect of all variants, or *a*_1_ = 1 and *a*_2_ = 25, which corresponds to upweighting more rare variants under the assumption that more rare variants have larger effects. The power of each test depends on the true disease model. However, the true disease model is unknown and variable in practice. To robustly aggregate information from different tests and weights, we propose an omnibus test using the Cauchy method via ACAT^28; 29^ and define the test statistic as

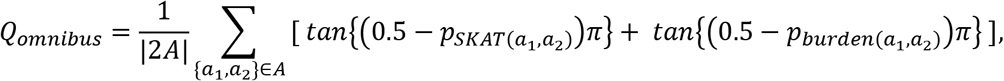

where *p*_*SKAT*(*a*_1_,*a*_2_)_ and *p*_*burden*(*a*_1_,*a*_2_)_ denote the p-value of SKAT and burden test using the beta function *beta*(*MAF, a*_1_, *a*_2_) = *MAF*^(*a*_1_−1)^(1 − *MAF*)^(*a*_2_−1)^ as the weight, and *A* is a set of specified values (*a*_1_,*a*_2_), and |*A*| is the size of set *A*. In the simulation studies, we set *A* = {(*a*_1_,*a*_2_)} = {(1,1), (1,25)}. Other choices of *a*_1_ and *a*_2_, e.g., (*a*_1_,*a*_2_) = (0.5,0.5) can also be used. The p-value of *Q_omnibus_* could be approximated by^29^

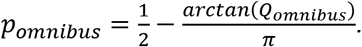

Compared to the minimum p-value combination, the Cauchy-based ACAT method is expected to increase power when more than one p-value of candidate tests are small. The proposed method also has two advantages over SKAT-O.^17^ First, it is flexible to the choices of weights, while SKAT-O is not able to combine SKAT and burden test under different choices of weights. Second, SKAT-O is computationally more expensive than the proposed omnibus test using ACAT.

#### Dynamic Detection of Rare Variants Association Regions with Different Window Sizes using Scan Statistic

Under the global null hypothesis, no variant is associated with a phenotype across the genome. Under the alternative hypothesis, there exist *r* signal regions that contain variants associated with the phenotype. We allow the variants-phenotype associated signal regions to have different sizes across the genome. Our goal is to detect whether any variants-phenotype associated region exists across the genome, and if they do exist, to identify the locations and sizes of these regions. Specifically, we first test the global null of variants-phenotype association region in the genome *H*_0_:*r* = 0, and if *H*_0_ is rejected, detect and estimate all the variants-phenotype associated regions.

SCANG tackles this hypothesis testing problem by using the extreme value of the set-based p-value of all candidate moving windows of different sizes in a range of windows of practical interest,

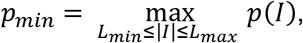

where *p*(*I*) is the p-value of region *I*, |*I*| denotes the number of variants in region *I*, and *L_min_* and *L_max_* are the smallest and largest variants number in windows of practical interest that are used for searching, respectively. For SCANG-S(*a*_1_,*a*_2_) and SCANG-B(*a*_1_,*a*_2_), *p*(*I*) is the p-value of SKAT and burden test using beta(MAF) weight with parameters *a*_1_ and *a*_2_, respectively. For SCANG-O, *p*(*I*) = *p_omnibus_* is the p-value of the proposed omnibus test which combines the information of burden test and SKAT using different weighting schemes using ACAT. A small value of *p_min_* indicates evidence against the null hypothesis. We reject the null hypothesis if the p-value of a region is smaller than a given threshold which controls the genome/family-wise type I error at the *α* level. If this results in only one region, the estimated signal region, i.e., the variants-phenotype associated region, is *Î* = argmin_*L*_*min*_≤|*I*|≤*L*_*max*__ *p*(*I*). If this results in multiple overlapping regions, we localize the signal region as the interval whose p-value is smaller than the threshold and achieves the local minimum in the sense that the p-value of that region is smaller than the regions that overlap with it.

#### Threshold for Controlling the Genome/Family-Wise Type I Error Rate

Since the candidate search windows in SCANG overlap with each other, the test statistics of these windows are highly correlated, and hence the standard Bonferroni correction for multiple testing adjustment is too conservative for SCANG and leads to power loss. Therefore, we propose an empirical threshold to control for the genome-wise type I error rate at the *α* level. The empirical threshold is calculated based on Monte Carlo simulations. For each simulated set *b*, We sample a set of *n* x 1 vectors 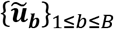 from a multivariate normal distribution *N*(0, ***I_n_***) and calculate the pseudo-score vector by 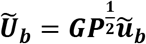, where 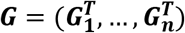 is the *p x n* genotype matrix, *p* is the total number of observed variants in the whole genome in a study and *n* is the number of subjects in the study, and *P* is the *n x n* projection matrix of the null GLM.^14; 30^ Note that every pseudo-score vector uses the same genotype matrix ***G*** and shares the same estimated *p x p* covariance matrix 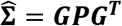. Then we calculate the region-based p-value of 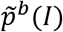 using the pseudo-score 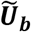. Although the individual score statistic might not be normally distributed for rare variants, the distributions of set-based test statistics are the same as those calculated using the pseudo-scores by assuming the normality for individual pseudo-scores, e.g., SKAT statistics using the observed scores and the pseudo-scores both follow a mixture of chi-square distribution. Hence the pseudo-p-value 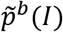 follows the same distribution as *p*(*I*). Finally we calculate the extreme value 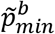 of 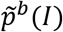 using the pseudo-score 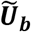. To control for the genome/family-wise type I error rate at the *α* level, we use the *α*th quantile of the empirical distribution of 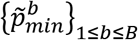 as the threshold. In practice, we select *B* = 2,000 for controlling the genome-wise/family-wise type I error rate, for example, at the 0.05 level.

#### Searching Algorithm for Multiple Signal Regions

In general, there might be several regions associated with the phenotype in the whole genome sequence. We note that the signal regions are short relatively to the size of the whole genome, and reasonably well separated in WGS. Hence intuitively, the scan statistic (p-value) of a properly estimated signal region should achieve a local minimum. On the other hand, the estimated signal region should also achieve genome-wise significance. We now describe an algorithm for detecting multiple signal regions (variants-phenotype association regions) as follows.

1. Set a genome-wise/family-wise significance level *α*, e.g, *α* = 0.05, the largest number of variants used in searching windows *L_max_*, the smallest number of variants used in searching windows *L_min_* and an overlap fraction 0 ≤ *f* ≤ 1, whose explanation is provided below. Calculate the empirical threshold *h*(*α, L_min_, L_max_*) for controlling the genome/family-wise type I error rate at the *α* level.
2. Calculate the *p*(*I*) for all the windows whose numbers of variants |*I*|’*s* are between *L_min_* and *L_max_*.
3. Pick the candidate set

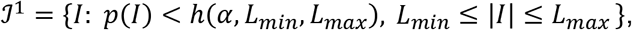

where *h*(*α, L_min_, L_max_*) is the empirical threshold to control for the genome-wise type I error at the *α* level. If 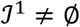, we reject the null hypothesis. Then rank the p-values of all the regions in the candidate set from the smallest to the largest, and set *j* = 1 and proceed with the following steps.
4. Let 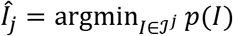, remove all the regions that overlap by more than the prespecified overlap fraction *f* with *Î_j_* from the candidate set, and update the candidate set as 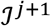, that is, 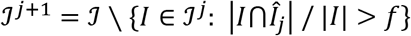.
5. Repeat step 4 with *j* = *j* + 1 until 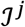 is an empty set.
6. Define *Î*_1_,*Î*_2_,… as the estimated signal regions.

**Figure S21** in the supplementary materials shows a simple example of this searching algorithm. Specification of the minimum and maximum searching window lengths *L_min_* and *L_max_* is an important issue in SCANG. Specifically, the range of the searching window length should be wide enough to ensure that each signal region will be searched. In the meantime, the range of the searching window length determines computation complexity. A smaller range of searching window lengths requires less computation. In practice, we choose the smallest and largest searching window lengths *L_min_* and *L_max_* based on sliding windows with multiple sizes, for example, in the simulation studies, we set *L_min_* as the 1%st percentile of the number of variants in all the 3kb sliding windows of simulated genome, and *L_max_* as the 99%th percentile of the number of variants in all 7 kb sliding windows of simulated genomes for the given sample size.

Since we consider all possible locations and multiple sizes of moving windows within the range *L_min_* ≤ |*I*| ≤ *L_max_* in the SCANG procedure, there might be multiple overlapped regions which achieve genome-wise significance. The fraction *f* is defined as the proportion of overlapped variants and controls for the degree of overlapping of detected regions. For example, when *f* = 0, the detected regions are non-overlapping with each other, and when *f* = 1, we keep every region below the genome-wise/family-wise threshold as the detected regions. In simulation studies and real data analysis, we allow the detected signal regions to be overlapping and set the overlap fraction *f* = 0.5. To reduce the computation time, we consider the searching window length with a skip of 5 variants.

### Simulation Studies

To validate SCANG in terms of protecting the genome/family-wise type I error and to assess its power compared to the conventional sliding window procedure, which considers sliding windows with a fixed length and skip length, we carried out simulation studies using a wide range of configurations. For all simulations, we generated sequence data by simulating 20,000 chromosomes for a 5 Mb region on the basis of the calibration coalescent model that mimics the linkage disequilibrium (LD) structure of samples of African American by using COSI.^31^ The simulation studies used the 5 Mb sequence to represent the whole genome and focused on low frequency and rare variants (minor allele frequency, MAF < 0.05).

#### Type I Error Simulations

To investigate whether SCANG preserves the desired genome/family-wise type I error rate, we simulated continuous phenotypes using the model:

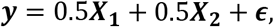

where ***X_x_*** is a continuous covariate generated from a standard normal distribution, ***X_2_*** is a dichotomous covariate taking values 0 and 1 with a probability of 0.5, and ***ϵ*** follows a standard normal distribution.

We repeated the type I error simulations for dichotomous phenotypes as above, except the dichotomous outcomes were generated via the model:

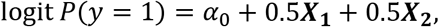

where ***X_1_*** and ***X_2_*** were continuous covariates, *α*_0_ was set to make the prevalence to 1%. Case-control sampling was used.

For both continuous and dichotomous simulations, we applied SCANG-B, SCANG-S and SCANG-O to 10^4^ replicates of genomes with size 5 MB, and examined the genome-wise type I error rate at α = 0.05 and 0.01. For SCANG-B and SCANG-S, we considered two weighting schemes, that is, for each variant *j, w_j_* = *Beta*(*MAF_j_*; 1,25) or *w_j_* = *Beta*(*MAF_j_*; 1,1) (unweighted). For SCANG-O, we used the ACAT method to construct the omnibus scan test by combining SKAT and burden in each window with two different weights *w_j_* = *Beta*(*MAF_j_*; 1,25) or *w_j_* = *Beta*(*MAF_j_*; 1,1) (unweighted) with the SCANG framework. The smallest and largest numbers of variants used in searching windows *L_min_* and *L_max_* were determined by the distributions of the numbers of variants in all sliding windows of 3 kb, 4 kb, 5 kb, 6kb and 7 kb in length. Specifically, we select the smallest number of variants of searching windows *L_min_* as the 1%st percentile of the number of variants in all the 3 kb sliding windows of simulated genomes, and the largest number of variants of searching windows *L_max_* as the 99%th percentile of the number of variants in all 7 kb sliding windows of simulated genomes. Note that *L_min_* and *L_max_* depend on the sample sizes n, which were set to be 2500, 5000 and 10000. To be specific, we set in the simulation *L_min_* = 45 and *L_max_* = 185 for *n* = 2,500, *L_min_* = 50 and *L_max_* = 200 for *n* = 5,000, and *L_min_ =* 70 and *L_max_* = 250 for *n* = 10,000.

#### Empirical Power Simulations

We randomly selected two signal regions (variants-phenotype association regions) across the 5 Mb sequence in each replicate. The length of the signal regions was randomly selected with lengths 3 kb, 4 kb, 5 kb and 6 kb. Within each signal region, 10% of variants were randomly chosen as causal variants with non-zero effect sizes *β*’s and the effect sizes of causal variants were set as a decreasing function of MAFs specified later in this paragraph. We generated continuous phenotypes by

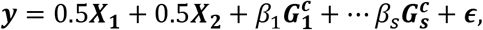

where ***X_1_, X_2_*** and ***ϵ*** are the same as those used in the type I error simulations, 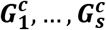 are the genotypes of the s causal variants in the two signal regions, and *β*s are effect sizes for causal variants. Similarly, we generated dichotomous phenotypes for case-control data under the logistic model

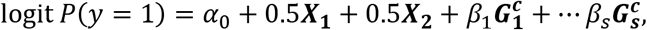

where *α*_0_, ***X_1_*** and ***X_2_*** are the same as those used in the type I error simulations, 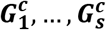 are again the genotypes of the s causal variants in the 2 signal regions, and *β*s are log ORs for the causal variants. For both models, we set the effect sizes as a decreasing function of MAFs *β_j_* = *c*|*log*_10_*MAF_j_*|, for continuous traits, *c* = 0.18, and for dichotomous traits, *c* = 0.255, which gives an OR of 3 for a variant with MAF = 5 x 10^−5^.

We applied SCANG-B, SCANG-S and SCANG-O to simulated datasets, and compared them with the corresponding conventional sliding window procedure where the sliding window size was set to be fixed at 3 kb, 4kb, 5kb, 6kb, 7 kb. For burden test and SKAT, we considered the same two weighting schemes as those in the type I error simulations, i.e., for the jth variant, *w_j_* = *Beta*(*MAF_j_*; 1,1) (unweighted) and *w_j_* = *Beta*(*MAF_j_*; 1,25).

We controlled the genome/family-wise type I error rate at the 0.05 level by using the proposed empirical threshold for SCANG and the Bonferroni correction for sliding window procedures. The ranges of the searching window lengths specified as the minimum and maximum numbers of variants in searching windows are the same as those used in the type I error rate simulation, i.e, (*L_min_, L_max_*) = (45,185), (50,200), (70,250) for sample sizes 2,500, 5,000, 10,000 respectively. Note that the lengths of signal regions are randomly set from 3 kb to 6 kb, hence the ranges of the searching window lengths specified using the smallest and the largest numbers of variants in 3 kb to 7 kb searching windows were larger than those of true signal regions. Burden test, SKAT and the proposed omnibus test were considered in the framework of SCANG and the conventional sliding window procedure.

In order to evaluate power, we considered two criteria. The first one is the causal variant detection rate, which is defined as

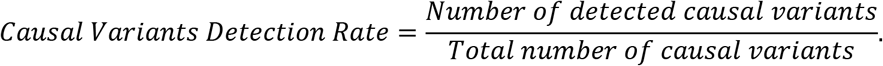

Here we defined a causal variant is detected if it is in one of detected signal regions. The causal variants detection rate could be regarded as the power of causal variants detection. The second one is the signal region detection rate, which is defined as

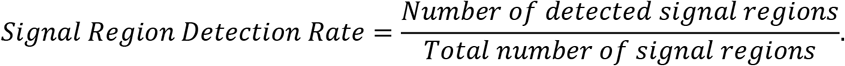

Here we defined the signal region as detected if it is overlapped with one of detected signal regions. Both the causal variant detection rate and the signal region detection rate can be regarded as the power of test.

### Application to ARIC WGS Data

The Atherosclerosis Risk in Communities (ARIC) study WGS data were generated by the Baylor College of Medicine Human Genome Sequencing Center. DNA samples were sequenced at 7.4-fold average depth on Illumina HiSeq instruments. After sample-level quality control,^20^ the ARIC WGS data consisted of around 55 million variants observed in 1,860 African Americans (AAs) and 33 million variants observed in 1,705 European Americans (EAs). Among all of the variants, 17.3% and 19.4% of them were common variants (MAF>5%) in AA and EA individuals, respectively. For this analysis, we focused on analyzing low frequency/rare variants across the genome, including low-frequency variants (1% ≤ MAF ≤ 5%, 13.4% in AA and 9.1% in EA) and rare variants (MAF < 1%, 69.3% in AA and 71.5% in EA).

To illustrate the proposed methods, we applied SCANG and a sliding window procedure for the whole-genome association analyses of two quantitative traits, sdLDL-c and Lp(a), both of which are related to cardiovascular disease risk.^20^ Following Morrison et.al,^20^ for the sliding window approach, we used the sliding window of length 4 kb and began at position 0 bp for each chromosome, with a skip length of 2 kb, and tested for the associations between the variants in each window and the phenotype using SKAT and burden test that weighted the variants by weights *w_j_* = *Beta*(*MAF*; 1,25) and no weight *w_j_* = *Beta*(*MAF*; 1,1), respectively. For the SCANG procedure, we applied the corresponding SCANG-S(1,25) and SCANG-B(1,1) to detect rare variant association regions. We further applied SCANG-O, which used the proposed omnibus test that aggregated SKAT(1,1), SKAT(1,25), Burden(1,1) and Burden(1,25) in the SCANG framework. We adjusted for age, sex and the first three principal components of ancestry in the analysis, consistent with that described in Morrison et al.^20^ Since the distribution of both sdLDL-c and Lp(a) are markedly skewed, we used the rank-based inverse normal transformed traits as phenotypes in the analysis.

We set the range of search window sizes in the SCANG procedures by specifying the minimum and maximum numbers of variants in searching windows between *L_min_* and *L_max_*, which was determined by the distributions of the numbers of variants in all the 3 kb, 4 kb, 5 kb, 6 kb and 7 kb sliding windows, that is, the 1%st percentile of the numbers of variants of all 3 kb sliding windows and the 99%th percentile of the numbers of variants of all 7 kb sliding windows. Since the number of variants in AAs is larger than that of EAs, *L_min_* and *L_max_* are different between AAs and EAs. Specifically, for AAs, *L_min_* = 20 and *L_max_* = 170, and for EAs, *L_min_* = 15 and *L_max_* = 120. We controlled the genome-wise/family-wise error rate (GWER/FWER) at the 0.05 level in the SCANG analysis using the proposed empirical threshold. For the sliding window procedure, followed Morrison et al,^20^ a minimum number of 3 minor allele counts were required in a window with a skip of length of 2kb, resulting in a total of 1,337,673 and 1,337,382 4 kb overlapping windows in AA and EA, respectively. As around 1.3 million windows were tested using the sliding window procedure, we used the Bonferroni method to control for the GWER/FWER at the 0.05 level. We hence set the region-based significance threshold for the sliding window procedure at 3.75 x 10^−8^ (approximately equal to 0.05/1,337,000). We note that SCANG directly controls for the GWER/FWER without the need of further multiple testing adjustment.

## Results

### Simulation of the Type I Error

The empirical genome-wise/family-wise type I error rates estimated for the SCANG methods, including SCANG-B(1,1), SCANG-B(1,25), SCANG-S(1,1), SCANG-S(1,25) and SCANG-O are presented in **Table 1** for α = 0.05 and 0.01 levels, respectively. All the five procedures have a well-controlled GWER type I error rate for both continuous and dichotomous traits at these significant levels, though for dichotomous traits in small sample size (n=2,500), SCANG-S(1,1) and SCANG-S(1,25) could be slightly conservative.

**Table 1.**
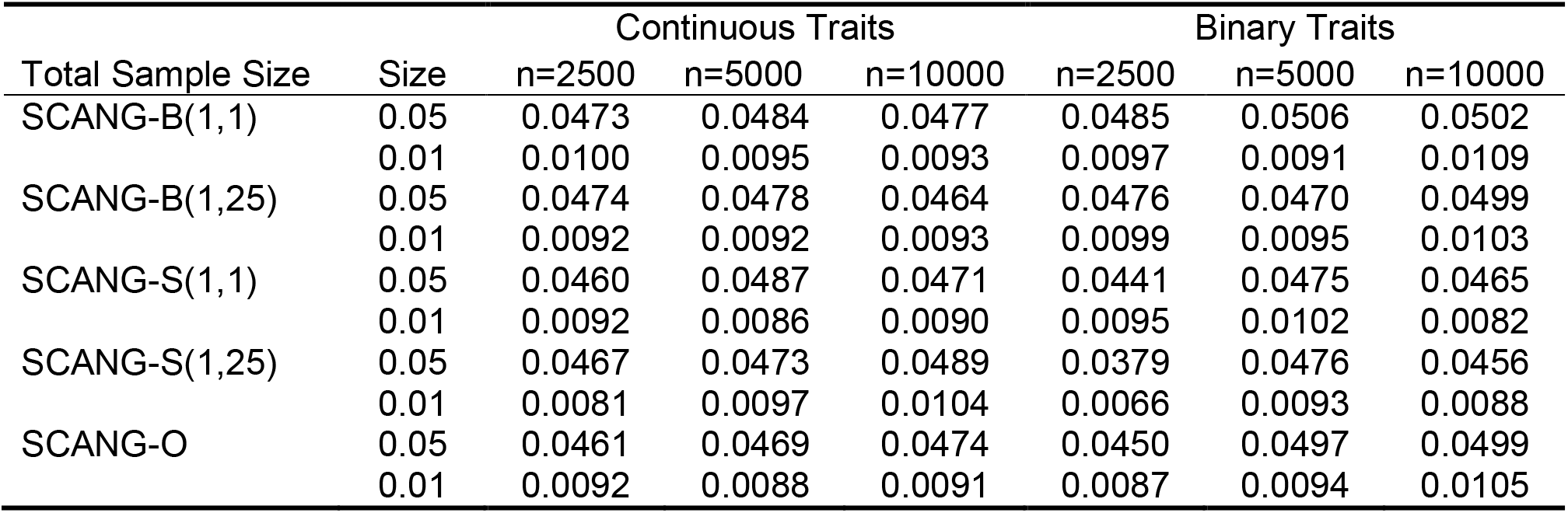
Genome-wise/family-wise empirical type I error rates from simulation studies using SCANG-B(1,1), SCANG-B(1,25), SCANG-S(1,1), SCANG-S(1,25) and SCANG-O at the genome-wide/family-wise significance level of α=0.05 and 0.01. SKAT(1,25) and Burden(1,1) are denoted by S(1,25) and B(1,1), and the corresponding SCANG methods are denoted by SCANG-S(1,25) and SCANG-B(1,1), the two numbers in the parentheses are the values (*a*_1_,*a*_2_) of the weight function beta(MAF, *a*_1_, *a_2_)*, The total sample size *n* was 2,500, 5,000 and 10,000. The smallest number of variants in searching windows was set as *L_min_* = 45, 50, 70 for *n* = 2,500, 5,000, 10,000, and the corresponding largest number of variants in searching windows was set as *L_max_* = 185, 200, 250. Each cell represents the empirical type I error rate estimate that was calculated as the proportion of p-values less than α under the null hypothesis based on 10^4^ replicates.

Additional results from simulations of the genome-wise type I error for SCANG using different ranges of searching window lengths are presented in **Table S1 and S2** for both continuous traits and dichotomous traits. They suggest that SCANG properly controls the genome-wise type I error rate at the nominal *α* = 0.05 or 0.01 levels for continuous traits. For dichotomous traits, SCANG-B(1,1), SCANG-B(1,25) and SCANG-O could also protect the genome-wise type I error, while SCANG-S(1,1) and SCANG-S(1,25) are conservative when the minimum searching length is small accompanying small sample sizes. In the current application, there were only include 20 variants in a region with a sample size of 2500, but it quickly approaches the nominal type I error rate 0.05 or 0.01 as the sample size increases.

#### Statistical Power of SCANG and Alternative Methods

We compared the power of SCANG with the conventional sliding window procedure in a series of simulation studies for both continuous and dichotomous traits by generating 5 Mb sequence data using a coalescent model. The ranges of the searching window lengths specified using the smallest and the largest numbers of variants in searching windows were much larger than those of true signal regions (**Figure S1**).

For continuous traits, SCANG had a much higher power than the sliding window procedure in the sense that SCANG had higher causal variants and signal region detection rates, and the advantages of SCANG were consistent for different choices of tests and weights (**Figure 1 and Figure S2**). For the sliding window procedures, the causal variants detection rate increased as the searching window length increased, while the signal region detection rate decreased as the searching window length increased. This indicates that, when the searching window length increases, the sliding window procedure tends to miss more signal regions, but the detected significant regions contain more causal variants. SCANG solves this problem by flexibly selecting the locations and the sizes of the signal regions.

**Figure 1.**
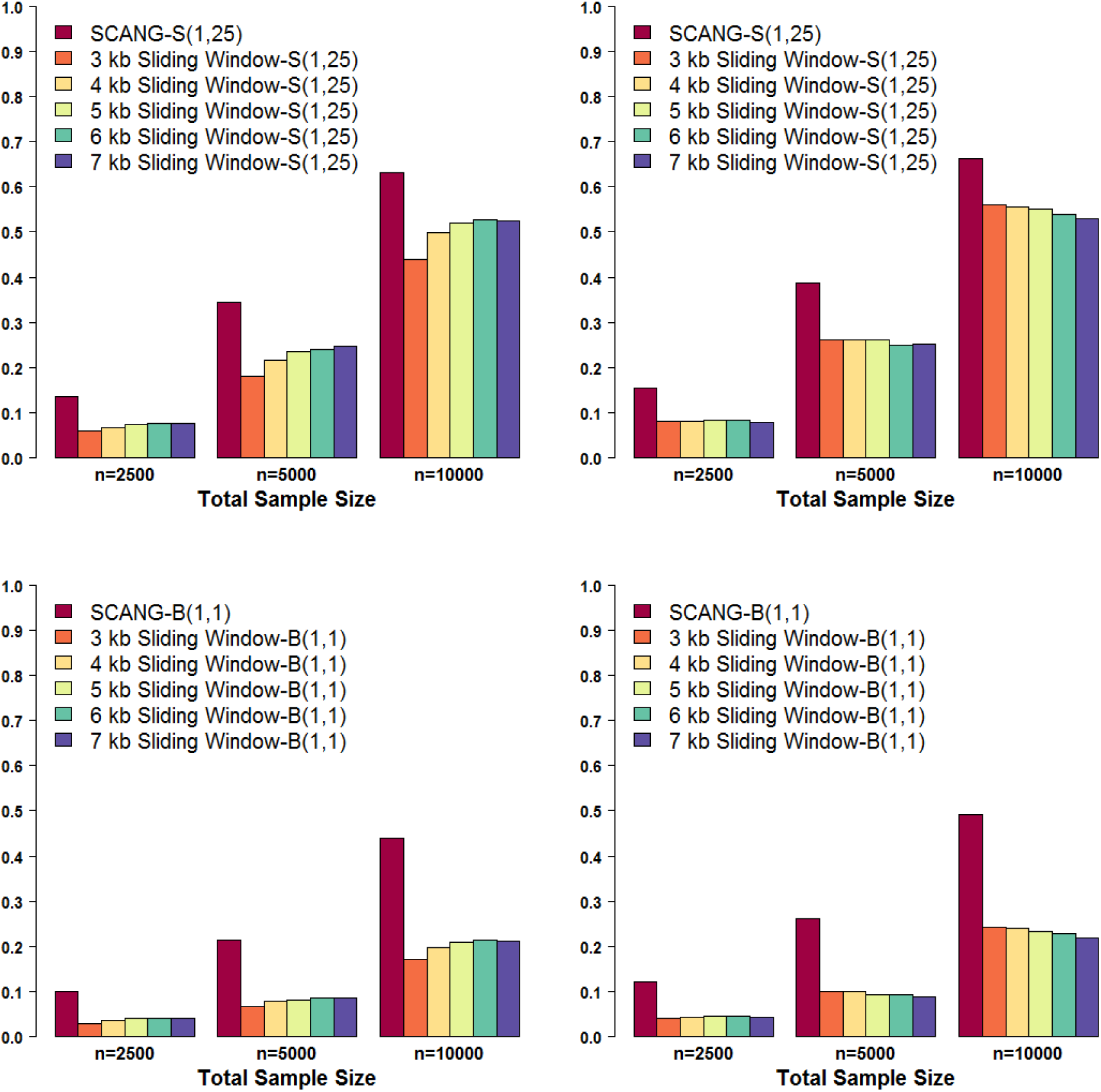
Power comparisons of SCANG and the corresponding sliding window procedures for continuous trait analysis. Empirical power was evaluated by the causal variants detection rate and the signal region detection rate defined in the simulation section. Both criteria were calculated at the genome-wise/family-wise type I error α = 0.05. We simulated 10% of the rare variants within random 3 kb-6 kb regions to be causal. We set for continuous phenotypes the maximum effect size equal to 0.774 for variants with MAF = 5 x 10^−5^ and let the effect sizes decrease with MAFs *β_j_* = *c*|*log*_10_*MAF_j_*|. The coefficients for the causal variants were all positive. The causal variants detection rate (left panel) is the proportion of detected causal variants in 2,000 simulated whole genome data sets, where a causal variant is called detected if it is in one of the detected signal regions. The signal region detection rate (right panel) is the proportion of detected signal regions in 2,000 simulated whole genome data sets, where a signal region is called detected if it is overlapped with one of the detected signal regions. For each configuration, the total sample sizes considered were 2,500, 5,000 and 10,000. For each setting, six methods were compared: SCANG and sliding window procedures using SKAT(1,25) and Burden(1,1), where the searching window lengths were equal to 3 kb, 4 kb, 5 kb, 6 kb and 7 kb. SKAT(1,25) and Burden(1,1) are denoted by S(1,25) and B(1,1), and the corresponding SCANG methods are denoted by SCANG-S(1,25) and SCANG-B(1,1), where the two numbers in the parentheses are the values (*a*_1_, *a*_2_) of the weight function beta(MAF, *a*_1_, *a*_2_). For SCANG, the range of search window lengths was set by the numbers of variants in searching windows between the 1%st percentile of the numbers of variants of all 3 kb sliding windows and the 99%th percentile of the numbers of variants of all 7 kb sliding windows.

The power difference between SCANG-S and the conventional sliding window method was similar using both weighting schemes, indicating that the substantial power gain of SCANG-S was robust to the choices of weights. Since only 10% of variants are causal variants in the signal regions, the power of SCANG-B was much lower than SCANG-S. However, the difference in power between SCANG-B and the conventional sliding window method was even larger, suggesting that the superior performance of SCANG is intrinsic and is not driven by the choices of set-based tests. The simulation results examining dichotomous traits were qualitatively similar and showed SCANG outperformed the conventional sliding window method, and the superior performance of SCANG was robust to the choices of set-based tests and weights (**Figure S3 and S4**).

We also conducted power simulations for different proportions of causal variants (40%) and different effect sizes in the signal regions. The results for both continuous and dichotomous phenotypes are presented in **Figures S5-S16**. They show that SCANG had much higher power than the conventional sliding window procedure in all considered settings. The power gain was also consistent for different choices of set-based tests and weights, and increased with the percentage of causal variants in signal regions.

Furthermore, with 10% causal variants in signal regions, the power of SCANG-S was higher than that of SCANG-B. With 40% causal variants in signal regions, the power of SCANG-B was higher than that of SCANG-S. In both scenarios, the power of SCANG-O was similar to the method that has the higher power (**Figure 2** and **Figure S17**), demonstrating that SCANG-O is robust and has omnibus power and could aggregate different association tests without losing much power.

**Figure 2.**
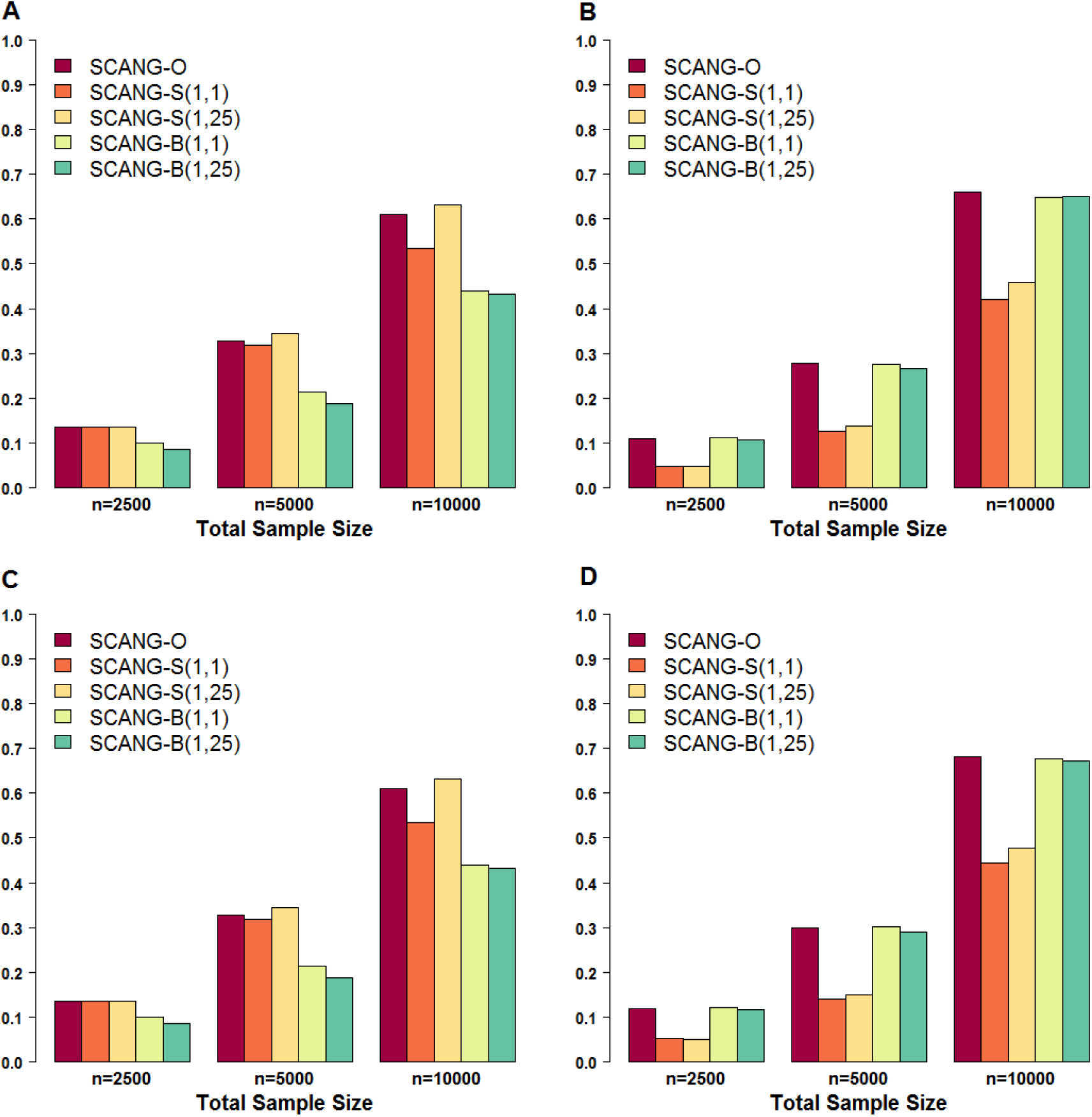
Power comparison of SCANG-O, SCANG-S(1,1), SCANG-S(1,25), SCANG-B(1,1) and SCANG-B(1,25) for continuous trait analysis. SKAT(*a*_1_, *a*_2_) and Burden(*a*_1_, *a*_2_) are denoted by S(*a*_1_, *a*_2_) and B(*a*_1_, *a*_2_), and the corresponding SCANG methods are denoted by SCANG-S(*a*_1_, *a*_2_) and SCANG-B(*a*_1_, *a*_2_). The two numbers in the parentheses are the values (*a*_1_, *a*_2_) of the weight function beta(MAF, *a*_1_, *a*_2_). Empirical power was evaluated by the causal variants detection rate and the signal region detection rate and both criteria were calculated at the genome-wise/family-wise type I error α = 0.05. We simulated 10% and 40% of the rare variants within random 3 kb-6 kb regions to be causal. We set for continuous phenotypes with the maximum effect size equal to 0.774 and 0.237 for variants with MAF = 5 x 10^−5^ when 10% and 40% causal variants, respectively, and let the effect sizes decrease with MAFs *β_j_* = *c*|*log*_10_*MAF_j_*|. The coefficients for the causal variants were all positive. The causal variants detection rate is the proportion of detected causal variants in 2,000 simulated data sets, where a causal variant is called detected if it is in one of the detected signal regions. The signal region detection rate is the proportion of detected signal regions in 2, simulated data sets, where a signal region is called detected if it is overlapped with one of the detected signal regions. The range of search window lengths was set by the numbers of variants in searching windows between the 1%st percentile of the numbers of variants of all 3 kb sliding windows and the 99%th percentile of the numbers of variants of all 7 kb sliding windows. For each configuration, the total sample sizes considered were 2,500, 5,000 and 10,000. (A) Causal variants detection rate when 10% causal variants. (B) Causal variants detection rate when 40% causal variants. (C) Signal region detection rate when 10% causal variants. (D) Signal region detection rate when 40% causal variants.

We further compared the power of SCANG-O with SCANG-S(1,1), SCANG-S(1,25), SCANG-B(1,1) and SCANG-B(1,25) for multiple effect sizes and different causal variants proportions (10% and 40%) of 10,000 samples, where the two numbers in the parentheses are the values (*a*_1_, *a*_2_) of the weight using the beta function beta(MAF, *a*_1_, *a*_2_). The results for continuous trait and binary trait are summarized in **Figure S18** and **Figure S19** in the Supplementary Data. The pattern was similar, and showed SCANG-O has a similar power to the method with the highest power if the genetic architecture were known, and is robust with little power loss by combining different tests.

#### Application to ARIC WGS Data

We compared the results using SCANG and the conventional sliding window procedure from the analysis of the lipid traits using the WGS data from the ARIC study. SCANG detected a region of 4,637 bp (from 45,382,398 to 45,387,034 bp on chromosome 19) consisting of 60 variants that had a significant association with sdLDL-c. Specifically, this detected region has a region-based p-value of 1.31 x 10^−9^ and genome-wise/family-wise p-value of 0.0445, among EAs using SCANG-O, which uses the ACAT method to combine SKAT(1,1), SKAT(1,25), Burden(1,1) and Burden(1,25) in each search window. This region resides in *NECTIN2* and covers two uncommon variants with individual variant-level p-values less than 1 x 10^−6^, including rs41290120 (*p* = 8.47 x 10^−9^), which is a pQTL of apolipoprotein E^32^, and rs283808 (*p* = 5.71 x 10^−7^). Note several common variants in *NECTIN2* have been found to have significant association with sdLDL-c in previous studies.^33; 34^ There is no significant region detected by SCANG-S(1,25) and SCANG-B(1,1). The significance of SCANG-O for this region was driven by SCANG-S(1,1).

The conventional 4 kb sliding window approaches used in the earlier analysis^20^ did not identify any genome-wide signal regions. Specifically, none of the 4 kb sliding windows covers both variants rs41290120 and rs283808 in *NECTIN2*. In the sliding windows that cover variant rs41290120, their p-values are much larger than that of the detected region from SCANG (**Table 2**). For example, the proposed omnibus test gives p-value 4.7 x 10^−8^ and 4.5 x 10^−8^ for the sliding windows indexed by rs41290120, which is larger than that of the detected region by SCANG-O (*p* = 1.3 x 10^−9^). The significant region detected by SCANG-O was evaluated for replication in ARIC AAs and was also strongly associated with sdLDL-c (*p* = 2.7 x 10^−4^), and was more significant than the sliding windows indexed by rs41290120 (**Table 3**).

**Table 2.** Set-based p-values of the significant regions detected by SCANG and two sliding windows indexed by variants rs41290120 on chromosome 19 in European Americans of the ARIC Whole Genome Sequencing Study (n=1,705) The two numbers in the parentheses are the The two numbers in the parentheses are the values (*a*_1_, *a*_2_) of the weight function beta(MAF, *a*_1_, *a*_2_). The physical positions of the windows are based on build hg19.

| Method | Start Pos.<br>(bp) | End Pos.<br>(bp) | Region<br>Size (bp) | SKAT<br>(1,25) | SKAT<br>(1,1) | Burden<br>(1,25) | Burden<br>(1,1) | Omnibus |
| --- | --- | --- | --- | --- | --- | --- | --- | --- |
| SCANG | 45,382,398 | 45,387,034 | 4,637 | $4.0 \times 10^{-5}$ | $3.3 \times 10^{-10}$ | $2.0 \times 10^{-2}$ | $1.3 \times 10^{-4}$ | $1.3 \times 10^{-9}$ |
| Sliding Window | 45,378,778 | 45,382,777 | 4,000 | $6.6 \times 10^{-6}$ | $1.2 \times 10^{-8}$ | $9.6 \times 10^{-1}$ | $1.3 \times 10^{-1}$ | $4.7 \times 10^{-8}$ |
| Sliding Window | 45,380,778 | 45,384,777 | 4,000 | $9.2 \times 10^{-7}$ | $1.2 \times 10^{-8}$ | $9.0 \times 10^{-2}$ | $8.3 \times 10^{-4}$ | $4.5 \times 10^{-8}$ |

**Table 3.** Replication of the significant regions detected by SCANG and two sliding windows indexed by variants rs41290120 on chromosome 19 in African Americans of the ARIC Whole Genome Sequencing Study (n=1,860) The two numbers in the parentheses are the The two numbers in the parentheses are the values (*a*_1_, *a*_2_) of the weight function beta(MAF, *a*_1_, *a*_2_). The physical positions of windows are based on build hg19.

| Method | Start Pos.<br>(bp) | End Pos.<br>(bp) | Region<br>Size (bp) | SKAT<br>(1,25) | SKAT<br>(1,1) | Burden<br>(1,25) | Burden<br>(1,1) | Omnibus |
| --- | --- | --- | --- | --- | --- | --- | --- | --- |
| SCANG | 45,382,398 | 45,387,034 | 4,637 | $1.2 \times 10^{-4}$ | $2.3 \times 10^{-4}$ | $2.2 \times 10^{-3}$ | $5.3 \times 10^{-4}$ | $2.7 \times 10^{-4}$ |
| Sliding Window | 45,378,778 | 45,382,777 | 4,000 | $1.6 \times 10^{-3}$ | $2.5 \times 10^{-3}$ | $1.4 \times 10^{-2}$ | $5.3 \times 10^{-3}$ | $3.1 \times 10^{-3}$ |
| Sliding Window | 45,380,778 | 45,384,777 | 4,000 | $5.0 \times 10^{-2}$ | $4.6 \times 10^{-2}$ | $9.1 \times 10^{-1}$ | $8.9 \times 10^{-1}$ | $1.6 \times 10^{-1}$ |

**Figure 3** and **Figure S20** summarize the genetic landscapes of the windows that are significantly associated with Lp(a) among AAs and EAs separately, using the five methods (SCANG-O, SCANG-S(1,25), SCANG-B(1,1) and the two sliding window procedures using SKAT(1,25) and Burden(1,1), where the two numbers in the parentheses are the values of beta(MAF) weight parameters *a*_1_ and *a*_2_, respectively. All of the significant windows resided in a 1.02 Mb region on chromosome 6 (from 160,501,196 bp to 161,524,329 bp), which includes seven genes *IGF2R* [MIM: 147280], *SLC22A1* [MIM: 602607], *SLC22A2* [MIM: 602608], *SLC22A3* [MIM: 604842], *LPAL2* [MIM: 611682], *LPA* [MIM: 152200], *PLG* [MIM: 173350] and *MAP3K4* [MIM: 602425], which have been previously shown to have common variants associated with Lp(a).^35–38^ Overall, the significant regions detected by SCANG-S(1,25) and SCANG-B(1,1) not only covered the significant windows detected by the corresponding sliding window procedures in both populations, but also detected several new regions that were missed by the sliding window method.

**Figure 3.**
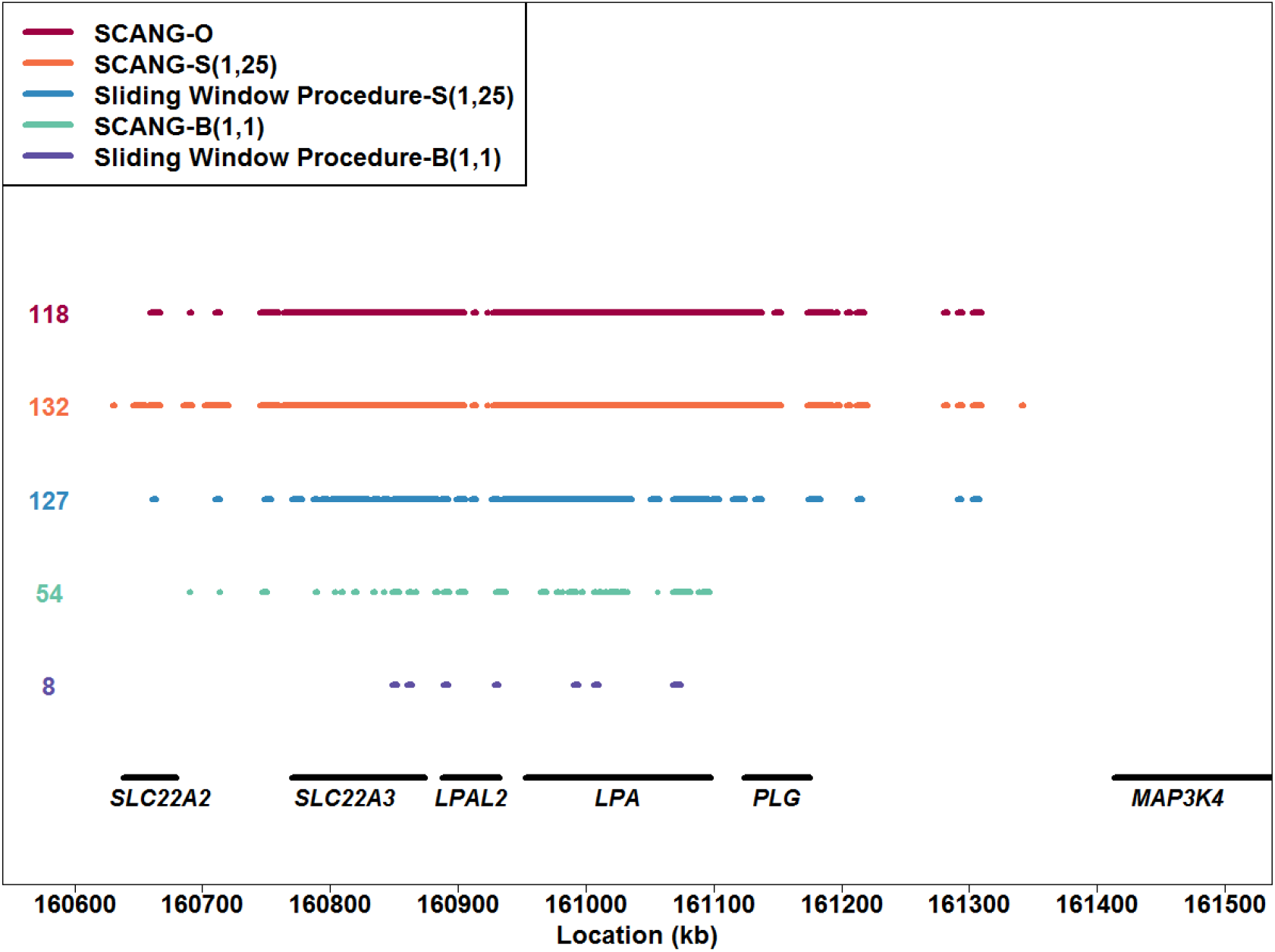
Genetic landscape of the windows significantly associated with Lp(a) levels on chromosome 6q25.3-6q26 among African Americans in the ARIC Whole Genome Sequencing Study (n=1,860) Five methods were compared: SCANG and 4kb sliding window procedures using SKAT(1,25) and Burden(1,1) (denoted by S(1,25) and B(1,1) respectively), with the corresponding SCANG methods denoted by SCANG-S(1,25) and SCANG-B(1,1), and SCANG-O, which used the proposed omnibus test that aggregated SKAT(1,1), SKAT(1,25), Burden(1,1) and Burden(1,25) using ACAT in the SCANG framework. The two numbers in the parentheses are the values (*a*_1_, *a*_2_) of the weight function beta(MAF, *a*_1_, *a*_2_), A dot means that the sliding window at this location is significant using the method that the color of the dot represents.

Specifically, SCANG-S(1,25) detected 132 and 64 significant regions, outperforming the 4 kb sliding window procedure with SKAT(1,25) which detected 127 and 35 significant sliding windows among AAs and EAs, respectively. SCANG-B(1,1) detected 54 and 19 significant regions, while the 4 kb sliding window procedure only detected 8 and 5 significant sliding windows among AAs and EAs, respectively. In addition, SCANG-S(1,25) and SCANG-B(1,1) were also able to detect several new regions which were separated from the regions detected by the corresponding sliding window procedures. For example, SCANG-S(1,25) could detect significant association between regions in *IGF2R* (from 160,501,196 bp to 160,507,026 bp) among EAs, which are at least 90 kb away from the detected sliding windows. The performance of SCANG-O was similar to SCANG-S(1,25), which detected most associations among the five methods. In summary, SCANG-O not only enables the identification of more significant findings, but also is less likely to miss important regions.

#### Computational Time

The computation time of SCANG depends on the sample size and the range of searching window lengths. To analyze a 5 Mb region sequenced on 2,500, 5,000 or 10,0 individuals, when we selected the searching window length between 3 kb and 7 kb, SCANG-O required 14 mins, 40 mins and 2 hours, respectively, on a 2.70 GHz laptop with 12 Gb memory. The computation cost decreased as the range of searching window length decreased, for example, it only required half of the time when we selected the searching window length between 3 kb and 5 kb. SCANG-O took 36 hours for analyzing the whole genome of EA individuals with ARIC whole genome sequencing data using the same single laptop. SCANG also works for parallel computing. Analyzing the whole genome on 10,000 individuals only requires 12 hours if using 100 computation cores for SCANG-O. Computing time can be further substantially reduced using cloud computing.^39^

## DISCUSSION

To overcome the limitations of the commonly used sliding window methods, which require pre-specifying fixed window lengths that are often unknown in practice, we propose SCANG, a dynamic and computationally efficient data adaptive scan procedure for use in WGS studies. SCANG is able to flexibly and powerfully detect rare variant associated regions in WGS by allowing the sizes of the variants-phenotype association regions to vary over the genome. SCANG allows for overlapping windows in scanning the genome and controls for the genome/family-wise error rate using an empirical threshold. Using extensive simulation studies, we demonstrate that SCANG is able to properly control for the genome/family-wise error rate, and the power of SCANG is greater than that of the conventional sliding window procedures using a fixed window size. Analysis of WGS and lipid data from the ARIC study illustrates the advantages of the proposed scan method for rare variants analysis. SCANG-O detected considerably more significant variants-phenotype association regions than the conventional sliding window method. For example, it was able to detect a significant association between the rare variants in a 4,637 bp region within *NECTIN2* and sdLDL-c among EA individuals that was missed by the sliding window procedure in the earlier analysis.^20^ These results show that SCANG improves the power of rare variants analysis by selecting the locations and the sizes of signal regions more precisely.

Instead of using fixed window sizes and skip lengths used in the conventional sliding window method, SCANG allows for multiple searching window sizes by considering all the moving windows of different sizes whose number of variants within a given range and allowing the searching windows to overlap. In our current numerical analysis, we set the range of sliding window sizes using the number of basepairs, e.g., 3 kb to 7 kb. As more rare variants will be observed in a given window with a fixed number of basepairs when the sample size increases, one can consider, for a larger sample size, a range of sliding window sizes using a smaller number of basepairs, e.g., 1kb to 7 kb, to detect smaller signal regions. Alternatively, one can specify a range of sliding window sizes using the number of variants, e.g., 40-200 variants as *L_min_* = 40 and *L_max_* = 200. Such a range specification using the number of variants is independent of sample sizes. Due to the flexibility of moving window selection, SCANG is able to identify the locations and the sizes of rare variant association regions from the data more accurately, and hence could dramatically increase the power of rare variant association analysis.

Different from some existing scan procedures, SCANG is a regression-based method that can be used for analyzing both continuous and discrete phenotypes. It allows for covariates adjustment, for example, ancestry PCs for population structure. By using the generalized linear mixed model framework, SCANG could also adjust for both population structure and relatedness. Specifically, when the data consist of related samples, following SMMAT,^40^ we consider the following Generalized Linear Mixed Model (GLMM)

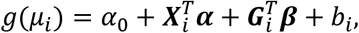

where the random effects *b_i_* account for population structure, relatedness and other types of between-observation correlation. We assume that 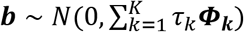 is an *n* x 1 vector of random effects *b_i_*, with the variance components *τ_k_* and the known *n x n* covariance matrices ***Φ_k_***. This means the random effects ***b*** can be decomposed into a sum of multiple random effects to account for different sources of relatedness and correlation as 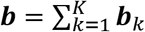 with ***b_k_*** ~ *N*(0, *τ_k_****Φ_k_***). For example, ***b***_1_ accounts for family relatedness with its covariance matrix ***Φ_1_*** as the Genetic Relatedness Matrix (GRM). Additional random effects (***b_2_, ⋯ b_K_***) can be used to account for complex sampling designs, such as hierarchical designs and correlation between repeated-measures from longitudinal studies using subject-specific random intercepts and slopes. After we fit the null model, we could proceed with SCANG by using the same steps as those described in the Method Section with the GLM scores replaced by the GLMM scores. SCANG could be easily extended to survival and multiple phenotypes framework and hence provides a comprehensive framework of WGS studies.

One attractive feature of the proposed method is that it allows for incorporating flexible weight functions to boost analysis power, for example, by giving larger weights to more rare variants and more functional variants which are more likely to be causal. Good choices of weights are likely to boost the power of SCANG. In our procedure, we consider continuous weights as a function of MAFs and give larger weights to more rare variants based on the fact that rare variants are more likely to be causal and variants with lower MAFs are likely to have larger effect sizes. However, this weight function also upweights neutral variants, for example, singletons and doubletons have the largest weights and most of them are neutral variants. One can also define weights using external bioinformatics and evolutionary biological knowledge by upweighting functional variants, such as using functional annotation information, e.g., protein scores, epigenetic scores, conservation scores, as well as integrative scores such as CADD^41^ and FATHMM-XF^42^.

The ability to obtain both individual region-level p-values without multiple comparison adjustment and their genome-wise/family-wise p-values with multiple comparison adjustment accounting for overlapping windows is another attractive feature of the proposed SCANG method. For each detected region, SCANG analytically calculates a variant set-based p-value, and also empirically calculates a genome-wise/family-wise p-value which adjusted for multiple testing. Note that the proposed procedure considers all possible regions across the genome and hence most of region-based p-values could be highly correlated. Instead of performing the usual Bonferroni correction, which could be quite conservative for multiple testing adjustment in this situation, we empirically calculate genome-wise/family-wise p-values by efficient Monte Carlo simulations. For each region, we calculate the region-based p-value using the score-based tests, such as burden and SKAT and the new omnibus tests that combine burden and SKAT and different weights using the Cauchy-based ACAT method.^29^ Calculations of these methods only require fitting the null model by regressing a phenotype on covariates once across the genome. Besides, we use the same genotype matrix ***G*** in each simulation and hence we only need to calculate the covariance matrix of the individual score test once. Therefore, the computational time of SCANG is affordable even with Monte Carlo simulations. For example, analyzing the entire genome on 10,000 individual requires 12 hours if we have 100 computation cores. However, calculating the empirical threshold still costs additional time. It is of future research interest to develop an analytic approximation to the genome-wide significance level for the proposed SCANG procedure.

### Supplemental Data

Supplemental Data include 21 figures and 2 tables.

## Acknowledgments

This work was supported by grants R35-CA197449, P01-CA134294, U01-HG009088, U19-CA203654, and R01-HL113338 (to X.L.). The Atherosclerosis Risk in Communities study has been funded in whole or in part with Federal funds from the National Heart, Lung, and Blood Institute, National Institutes of Health, Department of Health and Human Services (contract numbers HHSN268201700001I, HHSN268201700002I, HHSN268201700003I, HHSN268201700004I and HHSN268201700005I). The authors thank the staff and participants of the ARIC study for their important contributions. Funding support for “Building on GWAS for NHLBI-diseases: the U.S. CHARGE consortium” was provided by the NIH through the American Recovery and Reinvestment Act of 2009 (ARRA) (5RC2HL102419). Sequencing was carried out at the Baylor College of Medicine Human Genome Sequencing Center and supported by the National Human Genome Research Institute grants U54 HG003273 and UM1 HG008898.

## Web Resources

The URLs for software programs referenced in this article are as follows: SCANG, https://github.com/zilinli1988/SCANG and https://content.sph.harvard.edu/xlin/software.html Online Mendelian Inheritance in Man, http://www.omim.org

