## Supplemental Data for "Dynamic Scan Procedure for Detecting Rare-Variant Association Regions in Whole Genome Sequencing Studies"

**Figure S1. Number of variants in signal regions in simulation studies**

Physical length of signal regions in simulation studies was random selected from 3 kb, 4 kb, 5kb and 6 kb. For SCANG, variants number in searching windows was between 1%st quantile of variants number in 3 kb sliding windows and 99%th quantile of variants number in 7 kb sliding windows. $L_{min}$ and $L_{max}$ represents the smallest and largest variants number in the searching windows of SCANG. (A) Sample size n=2,500. (B) Sample size n=5,000. (C) Sample size n=10,000.

P

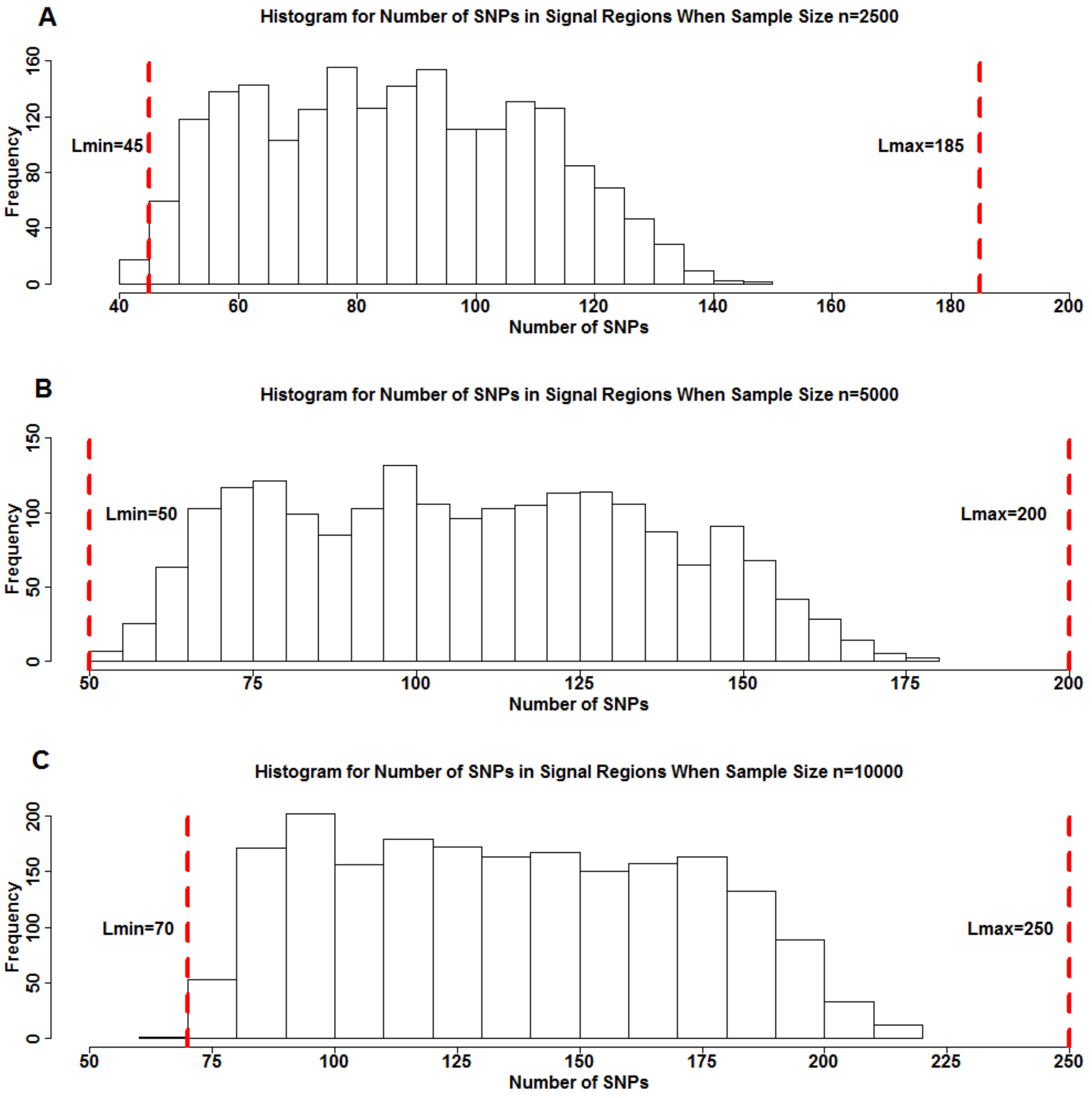

**Figure S2**. **Power comparisons of SCANG and the corresponding sliding window procedures using SKAT(1,1) and Burden(1,25) for continuous trait analysis when 10% of rare variants are causal variants**

Empirical power was evaluated by causal variants detection rate and signal region detection rate defined in the simulation section. Both two criteria were calculated at the genome-wise/family-wise type I error $\alpha = 0.05$. We simulated **10%** of the rare variants within random 3 kb-6 kb regions to be causal. We set for **continuous phenotypes** the maximum effect size equal to 0.774 for variants with $\mathrm{MAF} = 5 \times 10^{-5}$ and let the effect sizes decrease with MAFs $\beta_j = c|\, log_{10} MAF_j\,|$. The coefficients for the causal variants were all positive. The causal variants detection rate (left panel) is the proportion of detected causal variants in 2,000 simulated whole genome data sets, where a causal variant is called detected if it is in one of the detected signal regions. The signal region detection rate (right panel) is the proportion of detected signal regions in 2,000 simulated whole genome data sets, where a signal region is called detected if it is overlapped with one of the detected signal regions. For each configuration, the total sample size considered were 2,500, 5,000 and 10,000. For each setting, six methods were compared: SCANG and sliding window procedures using **SKAT(1,1)** and **Burden(1,25)** with searching window length used in the sliding window method equal to 3 kb, 4 kb, 5 kb, 6 kb and 7 kb. SKAT(1,1) and Burden(1,25) are denoted by S(1,1) and B(1,25), and the corresponding SCANG methods are denoted by SCANG-S(1,1) and SCANG-B(1,25) (the two numbers in the parentheses are the values of beta(MAF) weight parameters $a_1$ and $a_2$, respectively). For SCANG, the range of search window lengths was set by the numbers of variants in searching windows between the 1%st percentile of the numbers of variants of all 3 kb sliding windows and the 99%th percentile of the numbers of variants of all 7 kb sliding windows.

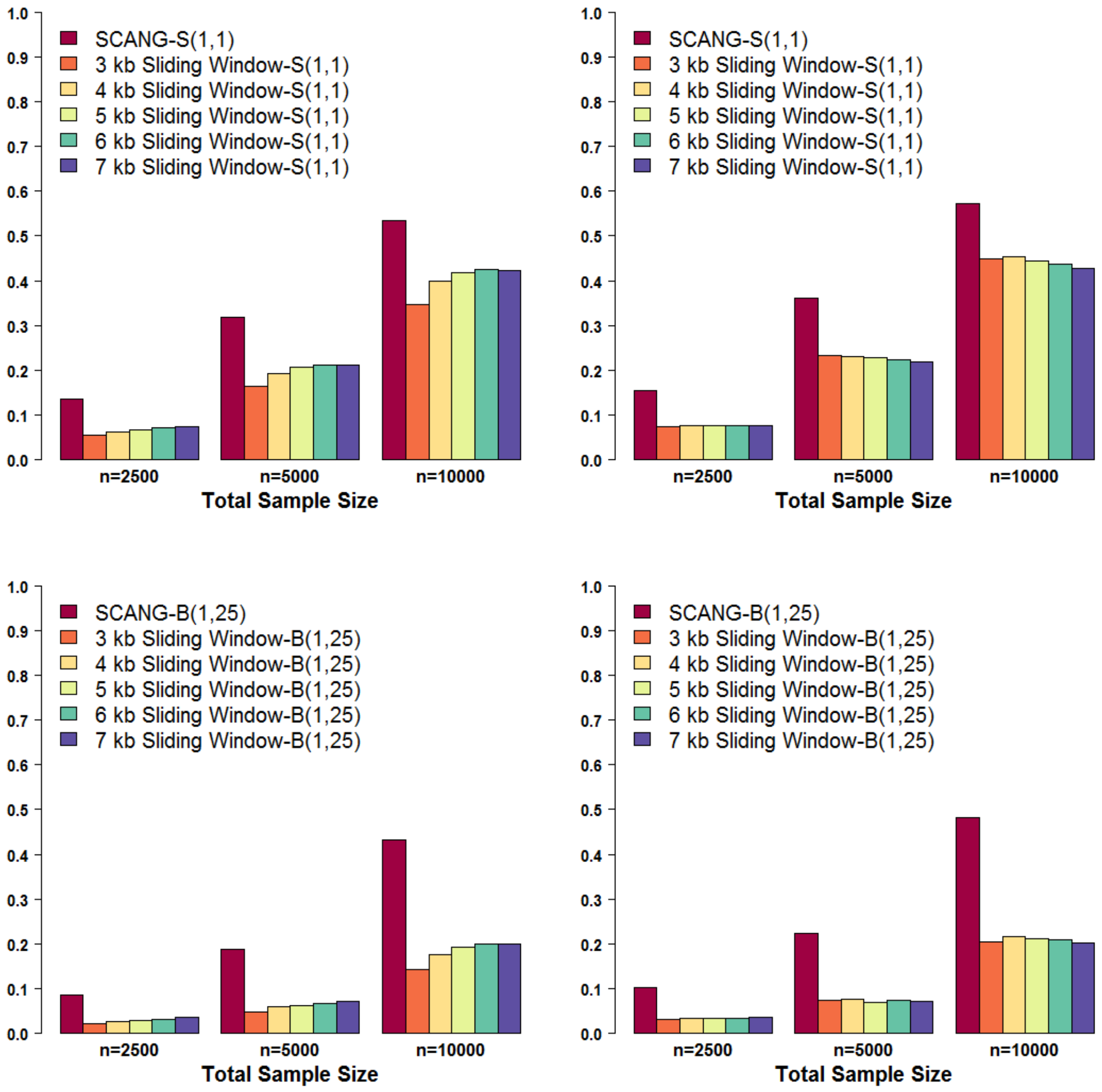

SCANG-S(1,1)
3 kb Sliding Window-S(1,1)
4 kb Sliding Window-S(1,1)
5 kb Sliding Window-S(1,1)
6 kb Sliding Window-S(1,1)
7 kb Sliding Window-S(1,1)
SCANG-B(1,25)
3 kb Sliding Window-B(1,25)
4 kb Sliding Window-B(1,25)
5 kb Sliding Window-B(1,25)
6 kb Sliding Window-B(1,25)
7 kb Sliding Window-B(1,25)
0.0
0.1
0.2
0.3
0.4
0.5
0.6
0.7
0.8
0.9
1.0
n=2500
n=5000
n=10000
Total Sample Size

**Figure S3. Power comparisons of SCANG and the corresponding sliding window procedures using SKAT(1,25) and Burden(1,1) for binary trait analysis when 10% of rare variants are causal variants**

Empirical power was evaluated by causal variants detection rate and signal region detection rate defined in the simulation section. Both two criteria were calculated at the genome-wise/family-wise type I error $\alpha = 0.05$. We simulated **10%** of the rare variants within random 3 kb-6 kb regions to be causal. We set for **binary phenotypes** with the maximum odds ratio equal to 3 for variants with $\mathrm{MAF} = 5 \times 10^{-5}$ and let the effect sizes decrease with MAFs $\beta_j = c| \log_{10} MAF_j |$. The coefficients for the causal variants were all positive. The causal variants detection rate (left panel) is the proportion of detected causal variants in 2,000 simulated whole genome data sets, where a causal variant is called detected if it is in one of the detected signal regions. The signal region detection rate (right panel) is the proportion of detected signal regions in 2,000 simulated whole genome data sets, where a signal region is called detected if it is overlapped with one of the detected signal regions. For each configuration, the total sample size considered were 2,500, 5,000 and 10,000. For each setting, six methods were compared: SCANG and sliding window procedures using **SKAT(1,25)** and **Burden(1,1)** with searching window length used in the sliding window method equal to 3 kb, 4 kb, 5 kb, 6 kb and 7 kb. SKAT(1,25) and Burden(1,1) are denoted by S(1,25) and B(1,1), and the corresponding SCANG methods are denoted by SCANG-S(1,25) and SCANG-B(1,1) (the two numbers in the parentheses are the values of beta(MAF) weight parameters $a_1$ and $a_2$, respectively). For SCANG, the range of search window lengths was set by the numbers of variants in searching windows between the 1%st percentile of the numbers of variants of all 3 kb sliding windows and the 99%th percentile of the numbers of variants of all 7 kb sliding windows.

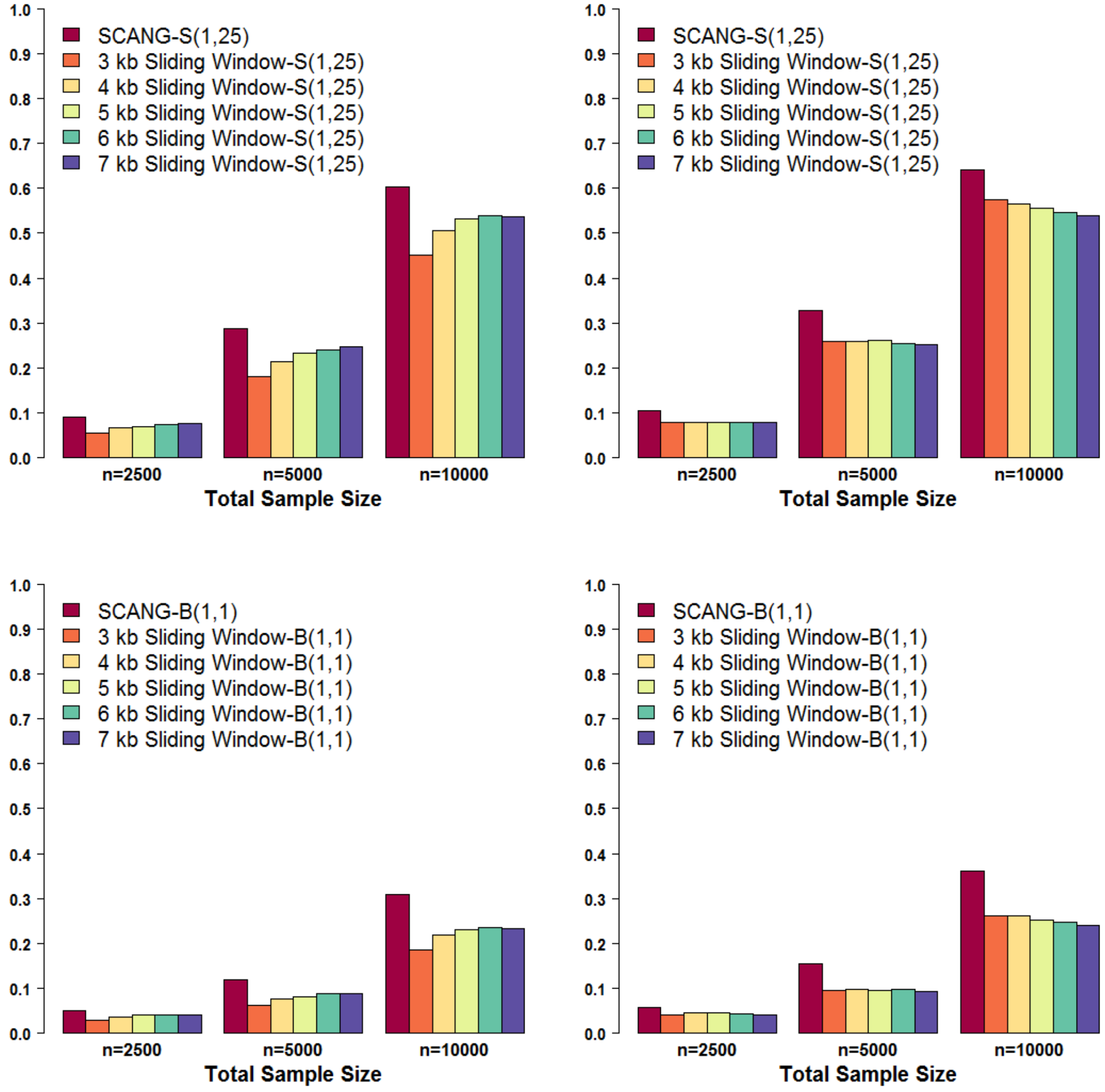

1.0
0.9
0.8
0.7
0.6
0.5
0.4
0.3
0.2
0.1
0.0
SCANG-S(1,25)
3 kb Sliding Window-S(1,25)
4 kb Sliding Window-S(1,25)
5 kb Sliding Window-S(1,25)
6 kb Sliding Window-S(1,25)
7 kb Sliding Window-S(1,25)
n=2500
n=5000
n=10000
Total Sample Size
SCANG-B(1,1)
3 kb Sliding Window-B(1,1)
4 kb Sliding Window-B(1,1)
5 kb Sliding Window-B(1,1)
6 kb Sliding Window-B(1,1)
7 kb Sliding Window-B(1,1)

**Figure S4**. **Power comparisons of SCANG and the corresponding sliding window procedures using SKAT(1,1) and Burden(1,25) for continuous trait analysis when 10% of rare variants are causal variants**

Empirical power was evaluated by causal variants detection rate and signal region detection rate defined in the simulation section. Both two criteria were calculated at the genome-wise/family-wise type I error $\alpha = 0.05$. We simulated **10%** of the rare variants within random 3 kb-6 kb regions to be causal. We set for **binary phenotypes** with the maximum odds ratio equal to 3 for variants with $\mathrm{MAF} = 5 \times 10^{-5}$ and let the effect sizes decrease with MAFs $\beta_j = c|\, log_{10} MAF_j \,|$. The coefficients for the causal variants were all positive. The causal variants detection rate (left panel) is the proportion of detected causal variants in 2,000 simulated whole genome data sets, where a causal variant is called detected if it is in one of the detected signal regions. The signal region detection rate (right panel) is the proportion of detected signal regions in 2,000 simulated whole genome data sets, where a signal region is called detected if it is overlapped with one of the detected signal regions. For each configuration, the total sample size considered were 2,500, 5,000 and 10,000. For each setting, six methods were compared: SCANG and sliding window procedures using **SKAT(1,1)** and **Burden(1,25)** with searching window length used in the sliding window method equal to 3 kb, 4 kb, 5 kb, 6 kb and 7 kb. SKAT(1,1) and Burden(1,25) are denoted by S(1,1) and B(1,25), and the corresponding SCANG methods are denoted by SCANG-S(1,1) and SCANG-B(1,25) (the two numbers in the parentheses are the values of beta(MAF) weight parameters $a_1$ and $a_2$, respectively). For SCANG, the range of search window lengths was set by the numbers of variants in searching windows between the 1%st percentile of the numbers of variants of all 3 kb sliding windows and the 99%th percentile of the numbers of variants of all 7 kb sliding windows.

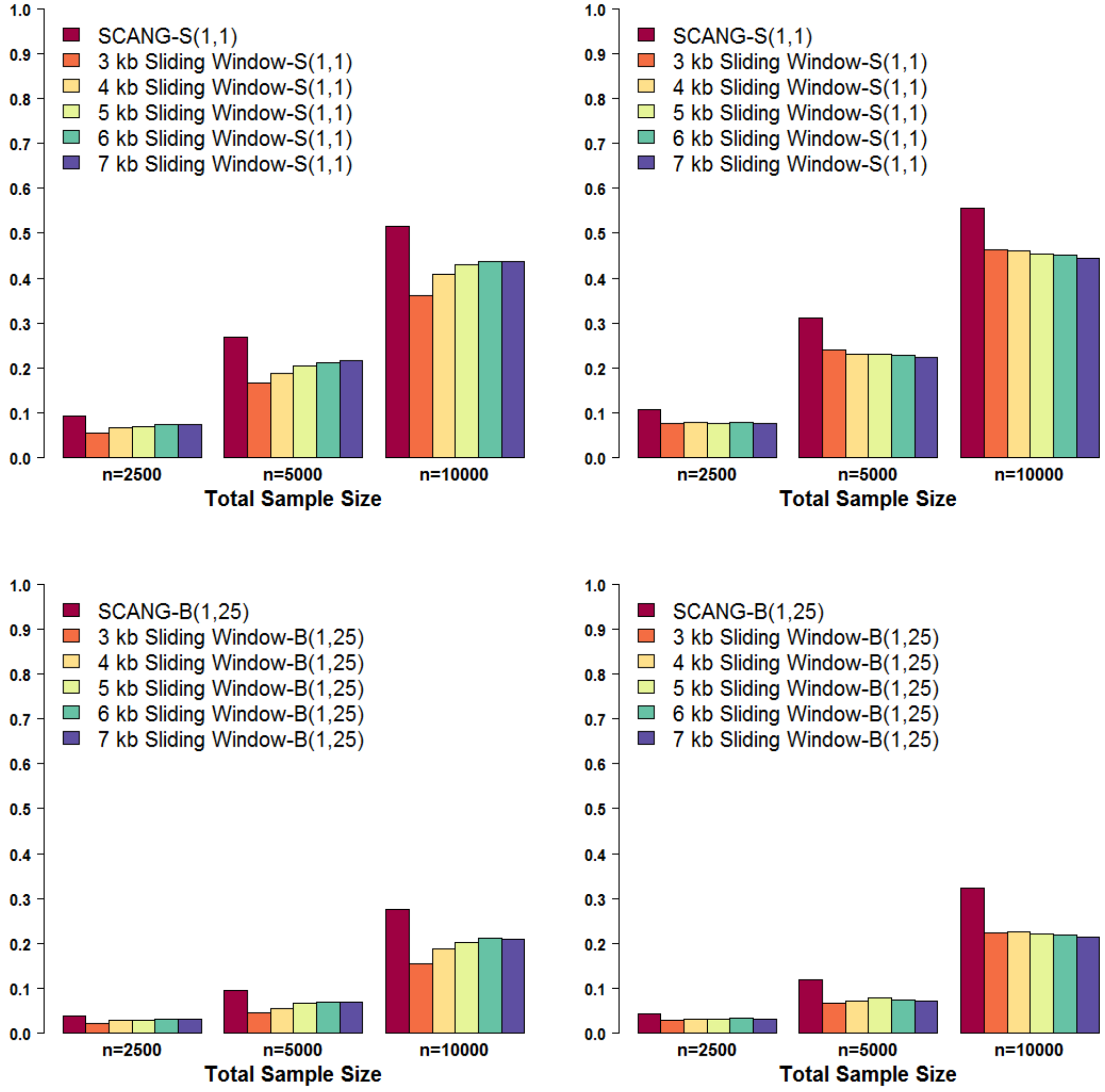
SCANG-S(1,1)
3 kb Sliding Window-S(1,1)
4 kb Sliding Window-S(1,1)
5 kb Sliding Window-S(1,1)
6 kb Sliding Window-S(1,1)
7 kb Sliding Window-S(1,1)
SCANG-B(1,25)
3 kb Sliding Window-B(1,25)
4 kb Sliding Window-B(1,25)
5 kb Sliding Window-B(1,25)
6 kb Sliding Window-B(1,25)
7 kb Sliding Window-B(1,25)
0.0
0.1
0.2
0.3
0.4
0.5
0.6
0.7
0.8
0.9
1.0
n=2500
n=5000
n=10000
Total Sample Size

**Figure S5**. **Power comparisons of SCANG and the corresponding sliding window procedures using SKAT(1,25) and Burden(1,1) for continuous trait analysis when 40% of rare variants are causal variants**

Empirical power was evaluated by causal variants detection rate and signal region detection rate defined in the simulation section. Both two criteria were calculated at the genome-wise/family-wise type I error $\alpha = 0.05$. We simulated **40%** of the rare variants within random 3 kb-6 kb regions to be causal. We set for **continuous phenotypes** the maximum effect size equal to 0.237 for variants with $\mathrm{MAF} = 5 \times 10^{-5}$ and let the effect sizes decrease with MAFs $\beta_j = c| log_{10} MAF_j |$. The coefficients for the causal variants were all positive. The causal variants detection rate (left panel) is the proportion of detected causal variants in 2,000 simulated whole genome data sets, where a causal variant is called detected if it is in one of the detected signal regions. The signal region detection rate (right panel) is the proportion of detected signal regions in 2,000 simulated whole genome data sets, where a signal region is called detected if it is overlapped with one of the detected signal regions. For each configuration, the total sample size considered were 2,500, 5,000 and 10,000. For each setting, six methods were compared: SCANG and sliding window procedures using **SKAT(1,25)** and **Burden(1,1)** with searching window length used in the sliding window method equal to 3 kb, 4 kb, 5 kb, 6 kb and 7 kb. SKAT(1,25) and Burden(1,1) are denoted by S(1,25) and B(1,1), and the corresponding SCANG methods are denoted by SCANG-S(1,25) and SCANG-B(1,1) (the two numbers in the parentheses are the values of beta(MAF) weight parameters $a_1$ and $a_2$, respectively). For SCANG, the range of search window lengths was set by the numbers of variants in searching windows between the 1%st percentile of the numbers of variants of all 3 kb sliding windows and the 99%th percentile of the numbers of variants of all 7 kb sliding windows.

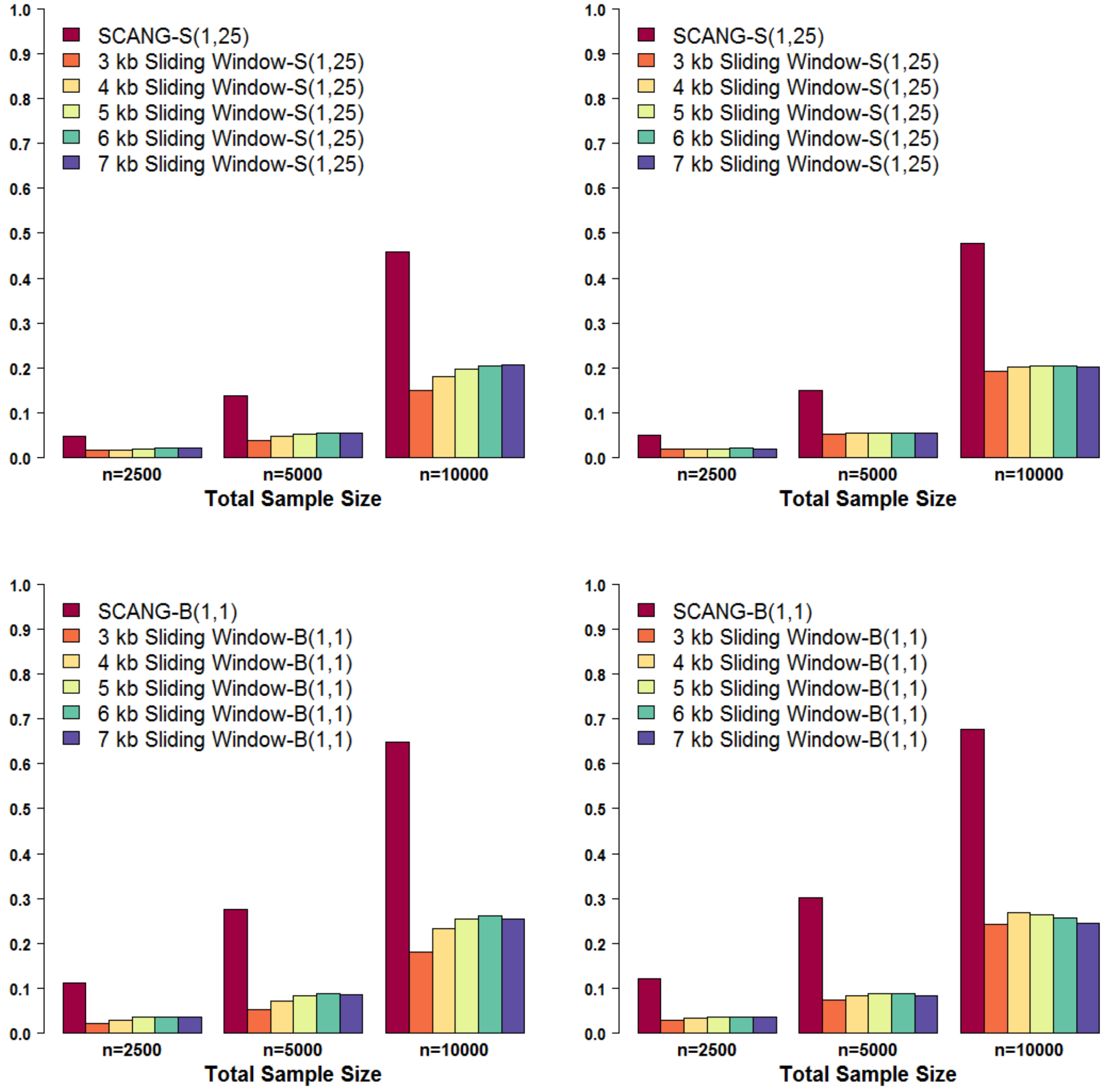

SCANG-S(1,25)
3 kb Sliding Window-S(1,25)
4 kb Sliding Window-S(1,25)
5 kb Sliding Window-S(1,25)
6 kb Sliding Window-S(1,25)
7 kb Sliding Window-S(1,25)
SCANG-B(1,1)
3 kb Sliding Window-B(1,1)
4 kb Sliding Window-B(1,1)
5 kb Sliding Window-B(1,1)
6 kb Sliding Window-B(1,1)
7 kb Sliding Window-B(1,1)
n=2500
n=5000
n=10000
Total Sample Size

**Figure S6**. **Power comparisons of SCANG and the corresponding sliding window procedures using SKAT(1,1) and Burden(1,25) for continuous trait analysis when 40% of rare variants are causal variants**

Empirical power was evaluated by causal variants detection rate and signal region detection rate defined in the simulation section. Both two criteria were calculated at the genome-wise/family-wise type I error $\alpha = 0.05$. We simulated **40%** of the rare variants within random 3 kb-6 kb regions to be causal. We set for **continuous phenotypes** the maximum effect size equal to 0.237 for variants with $\mathrm{MAF} = 5 \times 10^{-5}$ and let the effect sizes decrease with MAFs $\beta_j = c|\,log_{10} MAF_j\,|$. The coefficients for the causal variants were all positive. The causal variants detection rate (left panel) is the proportion of detected causal variants in 2,000 simulated whole genome data sets, where a causal variant is called detected if it is in one of the detected signal regions. The signal region detection rate (right panel) is the proportion of detected signal regions in 2,000 simulated whole genome data sets, where a signal region is called detected if it is overlapped with one of the detected signal regions. For each configuration, the total sample size considered were 2,500, 5,000 and 10,000. For each setting, six methods were compared: SCANG and sliding window procedures using **SKAT(1,1)** and **Burden(1,25)** with searching window length used in the sliding window method equal to 3 kb, 4 kb, 5 kb, 6 kb and 7 kb. SKAT(1,1) and Burden(1,25) are denoted by S(1,1) and B(1,25), and the corresponding SCANG methods are denoted by SCANG-S(1,1) and SCANG-B(1,25) (the two numbers in the parentheses are the values of beta(MAF) weight parameters $a_1$ and $a_2$, respectively). For SCANG, the range of search window lengths was set by the numbers of variants in searching windows between the 1%st percentile of the numbers of variants of all 3 kb sliding windows and the 99%th percentile of the numbers of variants of all 7 kb sliding windows.

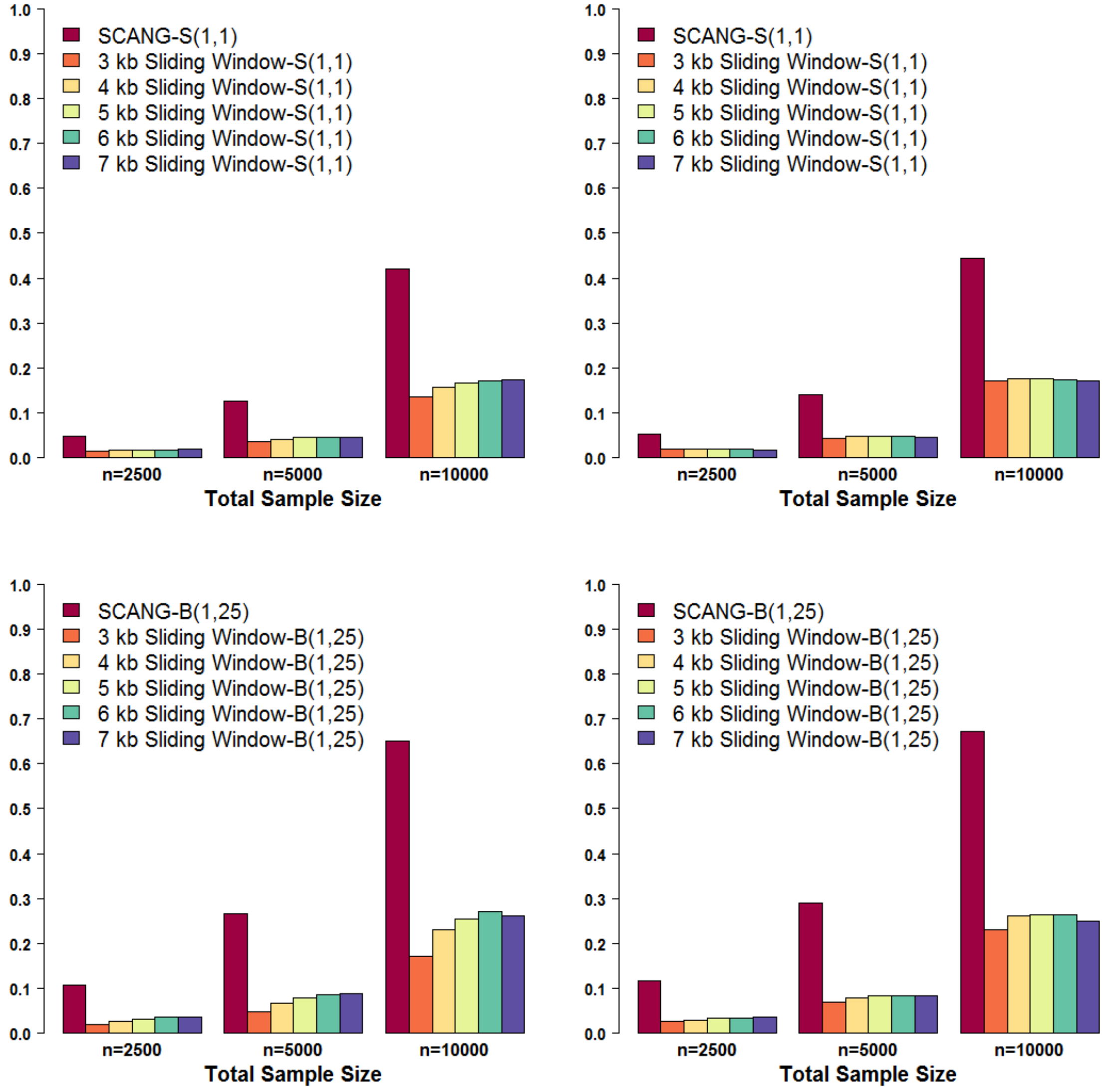

SCANG-S(1,1)
3 kb Sliding Window-S(1,1)
4 kb Sliding Window-S(1,1)
5 kb Sliding Window-S(1,1)
6 kb Sliding Window-S(1,1)
7 kb Sliding Window-S(1,1)
n=2500
n=5000
n=10000
Total Sample Size
SCANG-B(1,25)
3 kb Sliding Window-B(1,25)
4 kb Sliding Window-B(1,25)
5 kb Sliding Window-B(1,25)
6 kb Sliding Window-B(1,25)
7 kb Sliding Window-B(1,25)

**Figure S7. Power comparisons of SCANG and the corresponding sliding window procedures using SKAT(1,25) and Burden(1,1) for binary trait analysis when 40% of rare variants are causal variants**

Empirical power was evaluated by causal variants detection rate and signal region detection rate defined in the simulation section. Both two criteria were calculated at the genome-wise/family-wise type I error $\alpha = 0.05$. We simulated **40%** of the rare variants within random 3 kb-6 kb regions to be causal. We set for **binary phenotypes** with the maximum odds ratio equal to 1.65 for variants with $\mathrm{MAF} = 5 \times 10^{-5}$ and let the effect sizes decrease with MAFs $\beta_j = c|\, log_{10} MAF_j\,|$. The coefficients for the causal variants were all positive. The causal variants detection rate (left panel) is the proportion of detected causal variants in 2,000 simulated whole genome data sets, where a causal variant is called detected if it is in one of the detected signal regions. The signal region detection rate (right panel) is the proportion of detected signal regions in 2,000 simulated whole genome data sets, where a signal region is called detected if it is overlapped with one of the detected signal regions. For each configuration, the total sample size considered were 2,500, 5,000 and 10,000. For each setting, six methods were compared: SCANG and sliding window procedures using **SKAT(1,25)** and **Burden(1,1)** with searching window length used in the sliding window method equal to 3 kb, 4 kb, 5 kb, 6 kb and 7 kb. SKAT(1,25) and Burden(1,1) are denoted by S(1,25) and B(1,1), and the corresponding SCANG methods are denoted by SCANG-S(1,25) and SCANG-B(1,1) (the two numbers in the parentheses are the values of beta(MAF) weight parameters $a_1$ and $a_2$, respectively). For SCANG, the range of search window lengths was set by the numbers of variants in searching windows between the 1%st percentile of the numbers of variants of all 3 kb sliding windows and the 99%th percentile of the numbers of variants of all 7 kb sliding windows.

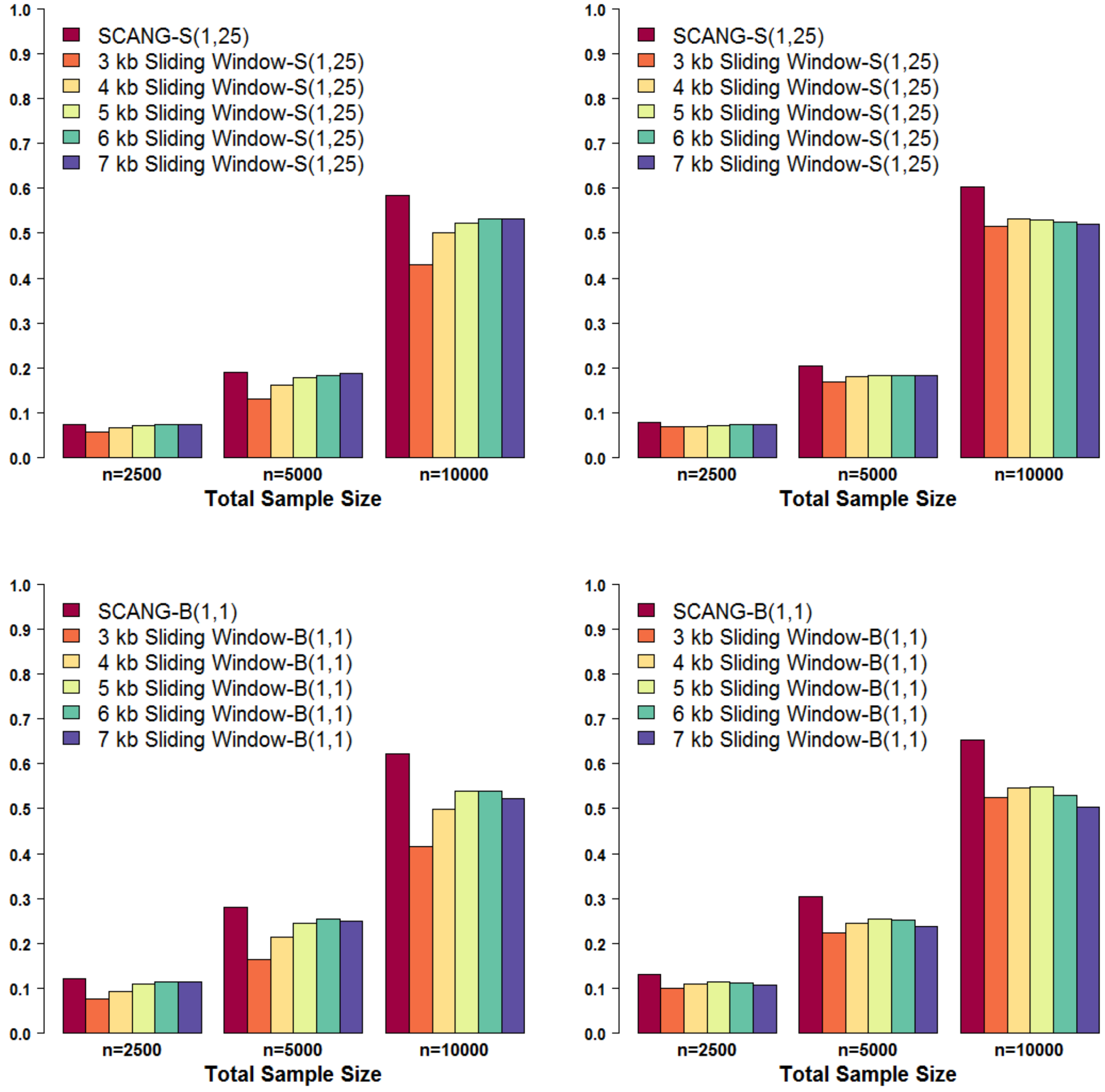
SCANG-S(1,25)
3 kb Sliding Window-S(1,25)
4 kb Sliding Window-S(1,25)
5 kb Sliding Window-S(1,25)
6 kb Sliding Window-S(1,25)
7 kb Sliding Window-S(1,25)
SCANG-B(1,1)
3 kb Sliding Window-B(1,1)
4 kb Sliding Window-B(1,1)
5 kb Sliding Window-B(1,1)
6 kb Sliding Window-B(1,1)
7 kb Sliding Window-B(1,1)
n=2500
n=5000
n=10000
Total Sample Size

**Figure S8**. **Power comparisons of SCANG and the corresponding sliding window procedures using SKAT(1,1) and Burden(1,25) for binary trait analysis when 40% of rare variants are causal variants**

Empirical power was evaluated by causal variants detection rate and signal region detection rate defined in the simulation section. Both two criteria were calculated at the genome-wise/family-wise type I error $\alpha = 0.05$. We simulated **40%** of the rare variants within random 3 kb-6 kb regions to be causal. We set for **binary phenotypes** with the maximum odds ratio equal to 1.65 for variants with $\mathrm{MAF} = 5 \times 10^{-5}$ and let the effect sizes decrease with MAFs $\beta_j = c|\, log_{10} MAF_j\,|$. The coefficients for the causal variants were all positive. The causal variants detection rate (left panel) is the proportion of detected causal variants in 2,000 simulated whole genome data sets, where a causal variant is called detected if it is in one of the detected signal regions. The signal region detection rate (right panel) is the proportion of detected signal regions in 2,000 simulated whole genome data sets, where a signal region is called detected if it is overlapped with one of the detected signal regions. For each configuration, the total sample size considered were 2,500, 5,000 and 10,000. For each setting, six methods were compared: SCANG and sliding window procedures using **SKAT(1,1)** and **Burden(1,25)** with searching window length used in the sliding window method equal to 3 kb, 4 kb, 5 kb, 6 kb and 7 kb. SKAT(1,1) and Burden(1,25) are denoted by S(1,1) and B(1,25), and the corresponding SCANG methods are denoted by SCANG-S(1,1) and SCANG-B(1,25) (the two numbers in the parentheses are the values of beta(MAF) weight parameters $a_1$ and $a_2$, respectively). For SCANG, the range of search window lengths was set by the numbers of variants in searching windows between the 1%st percentile of the numbers of variants of all 3 kb sliding windows and the 99%th percentile of the numbers of variants of all 7 kb sliding windows.

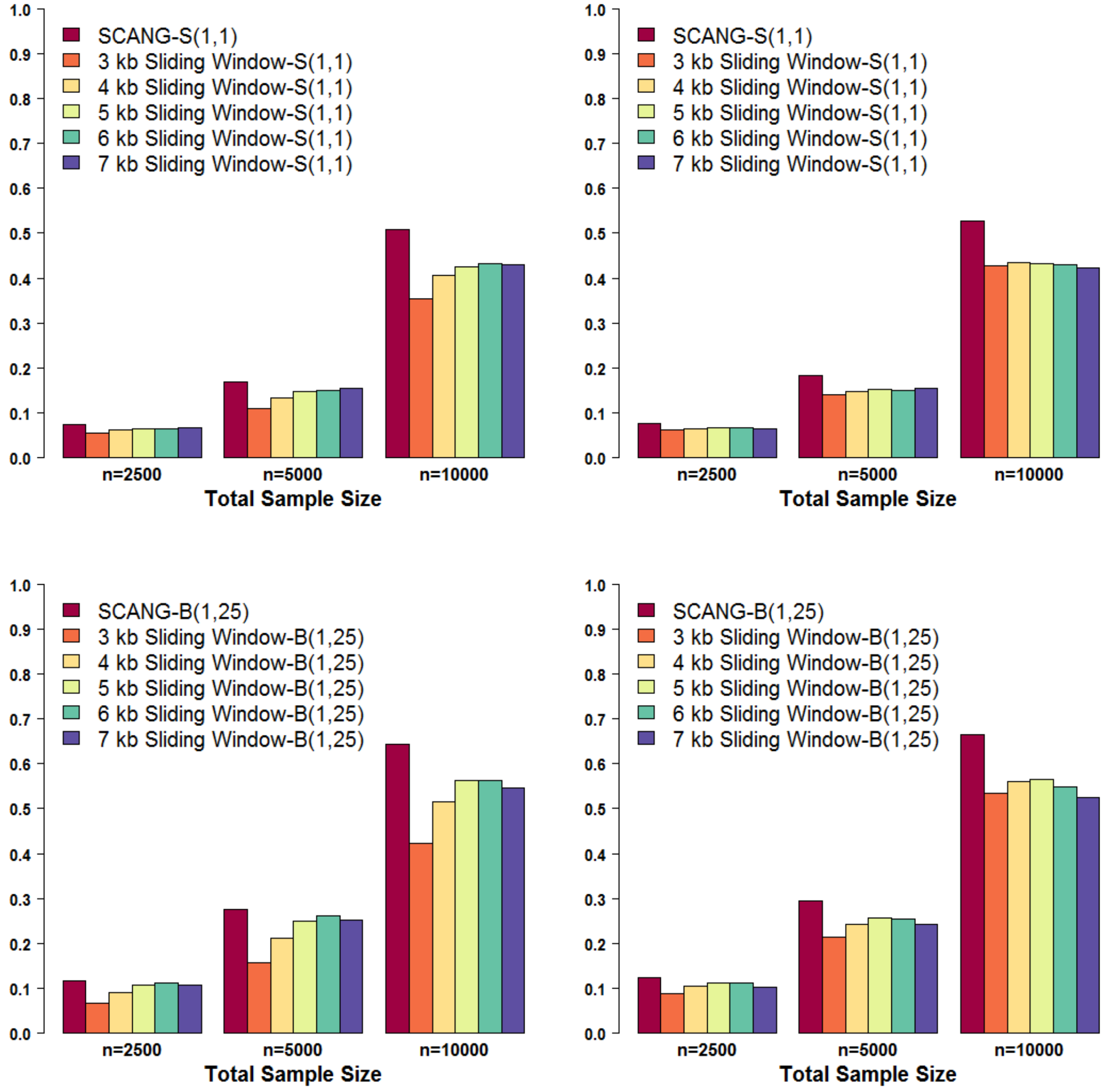
SCANG-S(1,1)
3 kb Sliding Window-S(1,1)
4 kb Sliding Window-S(1,1)
5 kb Sliding Window-S(1,1)
6 kb Sliding Window-S(1,1)
7 kb Sliding Window-S(1,1)
n=2500
n=5000
n=10000
Total Sample Size
SCANG-B(1,25)
3 kb Sliding Window-B(1,25)
4 kb Sliding Window-B(1,25)
5 kb Sliding Window-B(1,25)
6 kb Sliding Window-B(1,25)
7 kb Sliding Window-B(1,25)

**Figure S9**. **Power comparisons of SCANG and the corresponding sliding window procedures using SKAT(1,25) and Burden(1,1) for continuous trait analysis with multiple effect sizes when 10% of rare variants are causal variants**

Empirical power was evaluated by causal variants detection rate and signal region detection rate defined in the simulation section. Both two criteria were calculated at the genome-wise/family-wise type I error $\alpha = 0.05$. We simulated **10%** of the rare variants within random 3 kb-6 kb regions to be causal. We set for **continuous phenotypes** with the effect size $\beta_j = c|\,log_{10} MAF_j\,|$ where the effect sizes decrease with MAFs, and $c$ varies from 0.11 to 0.23. The coefficients for the causal variants were all positive. The causal variants detection rate (left panel) is the proportion of detected causal variants in 2,000 simulated whole genome data sets, where a causal variant is called detected if it is in one of the detected signal regions. The signal region detection rate (right panel) is the proportion of detected signal regions in 2,000 simulated whole genome data sets, where a signal region is called detected if it is overlapped with one of the detected signal regions. For each configuration, the total sample size considered were 10,000. For each setting, six methods were compared: SCANG and sliding window procedures using **SKAT(1,25)** and **Burden(1,1)** with searching window length used in the sliding window method equal to 3 kb, 4 kb, 5 kb, 6 kb and 7 kb. SKAT(1,25) and Burden(1,1) are denoted by S(1,25) and B(1,1), and the corresponding SCANG methods are denoted by SCANG-S(1,25) and SCANG-B(1,1) (the two numbers in the parentheses are the values of beta(MAF) weight parameters $a_1$ and $a_2$, respectively). For SCANG, the range of search window lengths was set by the numbers of variants in searching windows between the 1%st percentile of the numbers of variants of all 3 kb sliding windows and the 99%th percentile of the numbers of variants of all 7 kb sliding windows.

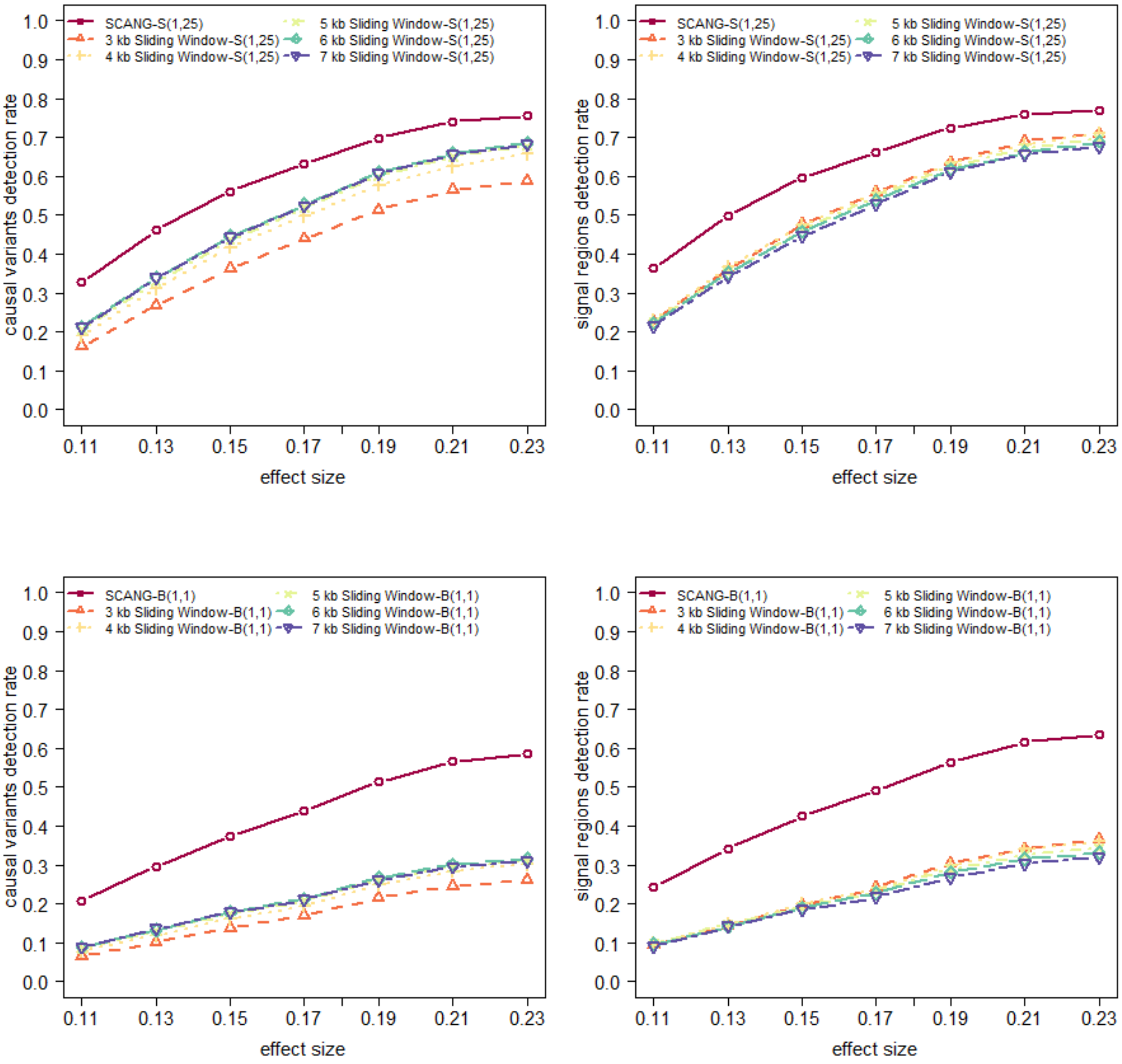
SCANG-S(1,25)
3 kb Sliding Window-S(1,25)
4 kb Sliding Window-S(1,25)
5 kb Sliding Window-S(1,25)
6 kb Sliding Window-S(1,25)
7 kb Sliding Window-S(1,25)
SCANG-B(1,1)
3 kb Sliding Window-B(1,1)
4 kb Sliding Window-B(1,1)
5 kb Sliding Window-B(1,1)
6 kb Sliding Window-B(1,1)
7 kb Sliding Window-B(1,1)
causal variants detection rate
signal regions detection rate
effect size
0.0 0.1 0.2 0.3 0.4 0.5 0.6 0.7 0.8 0.9 1.0
0.11 0.13 0.15 0.17 0.19 0.21 0.23

**Figure S10**. **Power comparisons of SCANG and the corresponding sliding window procedures using SKAT(1,1) and Burden(1,25) for continuous trait analysis with multiple effect sizes when 10% of rare variants are causal variants**

Empirical power was evaluated by causal variants detection rate and signal region detection rate defined in the simulation section. Both two criteria were calculated at the genome-wise/family-wise type I error $\alpha = 0.05$. We simulated **10%** of the rare variants within random 3 kb-6 kb regions to be causal. We set for **continuous phenotypes** with the effect size $\beta_j = c|\, log_{10} MAF_j\,|$ where the effect sizes decrease with MAFs, and $c$ varies from 0.11 to 0.23. The coefficients for the causal variants were all positive. The causal variants detection rate (left panel) is the proportion of detected causal variants in 2,000 simulated whole genome data sets, where a causal variant is called detected if it is in one of the detected signal regions. The signal region detection rate (right panel) is the proportion of detected signal regions in 2,000 simulated whole genome data sets, where a signal region is called detected if it is overlapped with one of the detected signal regions. For each configuration, the total sample size considered were 10,000. For each setting, six methods were compared: SCANG and sliding window procedures using **SKAT(1,1)** and **Burden(1,25)** with searching window length used in the sliding window method equal to 3 kb, 4 kb, 5 kb, 6 kb and 7 kb. SKAT(1,1) and Burden(1,25) are denoted by S(1,1) and B(1,25), and the corresponding SCANG methods are denoted by SCANG-S(1,1) and SCANG-B(1,25) (the two numbers in the parentheses are the values of beta(MAF) weight parameters $a_1$ and $a_2$, respectively). For SCANG, the range of search window lengths was set by the numbers of variants in searching windows between the 1%st percentile of the numbers of variants of all 3 kb sliding windows and the 99%th percentile of the numbers of variants of all 7 kb sliding windows.

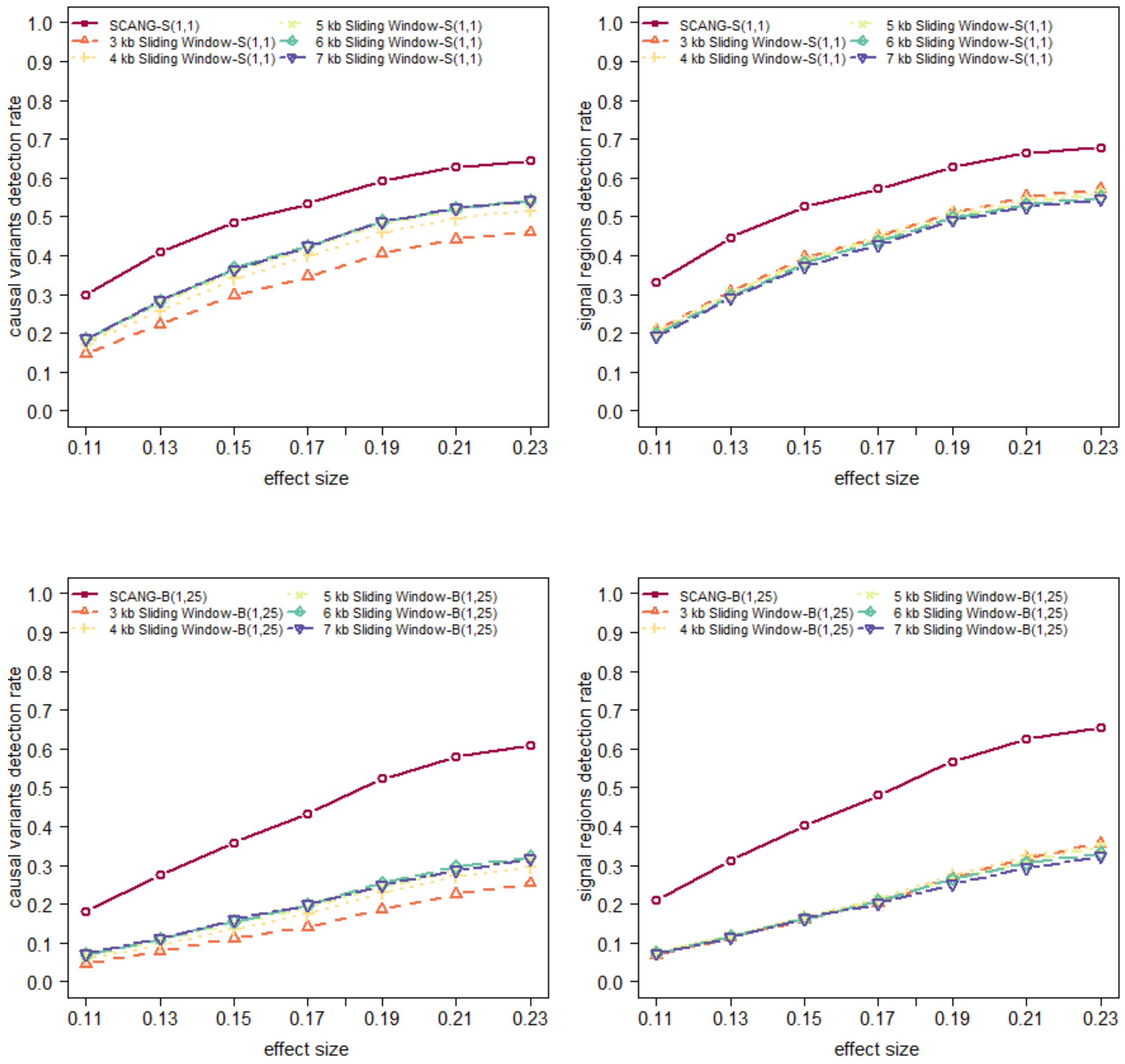

SCANG-S(1,1)
3 kb Sliding Window-S(1,1)
4 kb Sliding Window-S(1,1)
5 kb Sliding Window-S(1,1)
6 kb Sliding Window-S(1,1)
7 kb Sliding Window-S(1,1)
causal variants detection rate
signal regions detection rate
effect size
0.0
0.1
0.2
0.3
0.4
0.5
0.6
0.7
0.8
0.9
1.0
0.11
0.13
0.15
0.17
0.19
0.21
0.23
SCANG-B(1,25)
3 kb Sliding Window-B(1,25)
4 kb Sliding Window-B(1,25)
5 kb Sliding Window-B(1,25)
6 kb Sliding Window-B(1,25)
7 kb Sliding Window-B(1,25)

**Figure S11**. **Power comparisons of SCANG and the corresponding sliding window procedures using SKAT(1,25) and Burden(1,1) for continuous trait analysis with multiple effect sizes when 40% of rare variants are causal variants**

Empirical power was evaluated by causal variants detection rate and signal region detection rate defined in the simulation section. Both two criteria were calculated at the genome-wise/family-wise type I error $\alpha = 0.05$. We simulated **40%** of the rare variants within random 3 kb-6 kb regions to be causal. We set for **continuous phenotypes** with the effect size $\beta_j = c|\,log_{10} MAF_j\,|$ where the effect sizes decrease with MAFs, and $c$ varies from 0.04 to 0.07. The coefficients for the causal variants were all positive. The causal variants detection rate (left panel) is the proportion of detected causal variants in 2,000 simulated whole genome data sets, where a causal variant is called detected if it is in one of the detected signal regions. The signal region detection rate (right panel) is the proportion of detected signal regions in 2,000 simulated whole genome data sets, where a signal region is called detected if it is overlapped with one of the detected signal regions. For each configuration, the total sample size considered were 10,000. For each setting, six methods were compared: SCANG and sliding window procedures using **SKAT(1,25)** and **Burden(1,1)** with searching window length used in the sliding window method equal to 3 kb, 4 kb, 5 kb, 6 kb and 7 kb. SKAT(1,25) and Burden(1,1) are denoted by S(1,25) and B(1,1), and the corresponding SCANG methods are denoted by SCANG-S(1,25) and SCANG-B(1,1) (the two numbers in the parentheses are the values of beta(MAF) weight parameters $a_1$ and $a_2$, respectively). For SCANG, the range of search window lengths was set by the numbers of variants in searching windows between the 1%st percentile of the numbers of variants of all 3 kb sliding windows and the 99%th percentile of the numbers of variants of all 7 kb sliding windows.

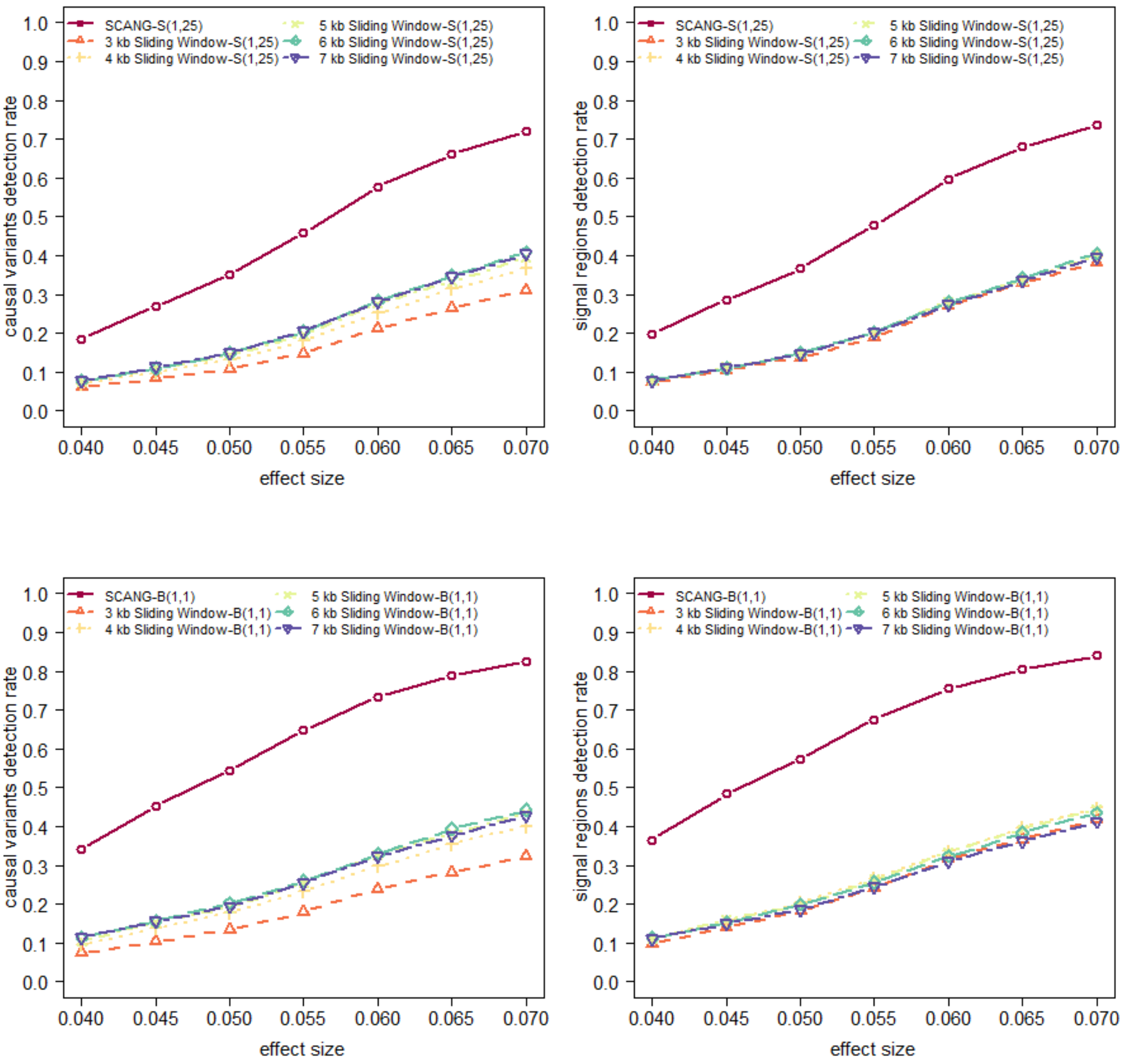
SCANG-S(1,25)
3 kb Sliding Window-S(1,25)
4 kb Sliding Window-S(1,25)
5 kb Sliding Window-S(1,25)
6 kb Sliding Window-S(1,25)
7 kb Sliding Window-S(1,25)
causal variants detection rate
signal regions detection rate
effect size
0.0
0.1
0.2
0.3
0.4
0.5
0.6
0.7
0.8
0.9
1.0
0.040
0.045
0.050
0.055
0.060
0.065
0.070
SCANG-B(1,1)
3 kb Sliding Window-B(1,1)
4 kb Sliding Window-B(1,1)
5 kb Sliding Window-B(1,1)
6 kb Sliding Window-B(1,1)
7 kb Sliding Window-B(1,1)

**Figure S12**. **Power comparisons of SCANG and the corresponding sliding window procedures using SKAT(1,1) and Burden(1,25) for continuous trait analysis with multiple effect sizes when 40% of rare variants are causal variants**

Empirical power was evaluated by causal variants detection rate and signal region detection rate defined in the simulation section. Both two criteria were calculated at the genome-wise/family-wise type I error $\alpha = 0.05$. We simulated **40%** of the rare variants within random 3 kb-6 kb regions to be causal. We set for **continuous phenotypes** with the effect size $\beta_j = c|\, log_{10} MAF_j\,|$ where the effect sizes decrease with MAFs, and $c$ varies from 0.04 to 0.07. The coefficients for the causal variants were all positive. The causal variants detection rate (left panel) is the proportion of detected causal variants in 2,000 simulated whole genome data sets, where a causal variant is called detected if it is in one of the detected signal regions. The signal region detection rate (right panel) is the proportion of detected signal regions in 2,000 simulated whole genome data sets, where a signal region is called detected if it is overlapped with one of the detected signal regions. For each configuration, the total sample size considered were 10,000. For each setting, six methods were compared: SCANG and sliding window procedures using **SKAT(1,1)** and **Burden(1,25)** with searching window length used in the sliding window method equal to 3 kb, 4 kb, 5 kb, 6 kb and 7 kb. SKAT(1,1) and Burden(1,25) are denoted by S(1,1) and B(1,25), and the corresponding SCANG methods are denoted by SCANG-S(1,1) and SCANG-B(1,25) (the two numbers in the parentheses are the values of beta(MAF) weight parameters $a_1$ and $a_2$, respectively). For SCANG, the range of search window lengths was set by the numbers of variants in searching windows between the 1%st percentile of the numbers of variants of all 3 kb sliding windows and the 99%th percentile of the numbers of variants of all 7 kb sliding windows.

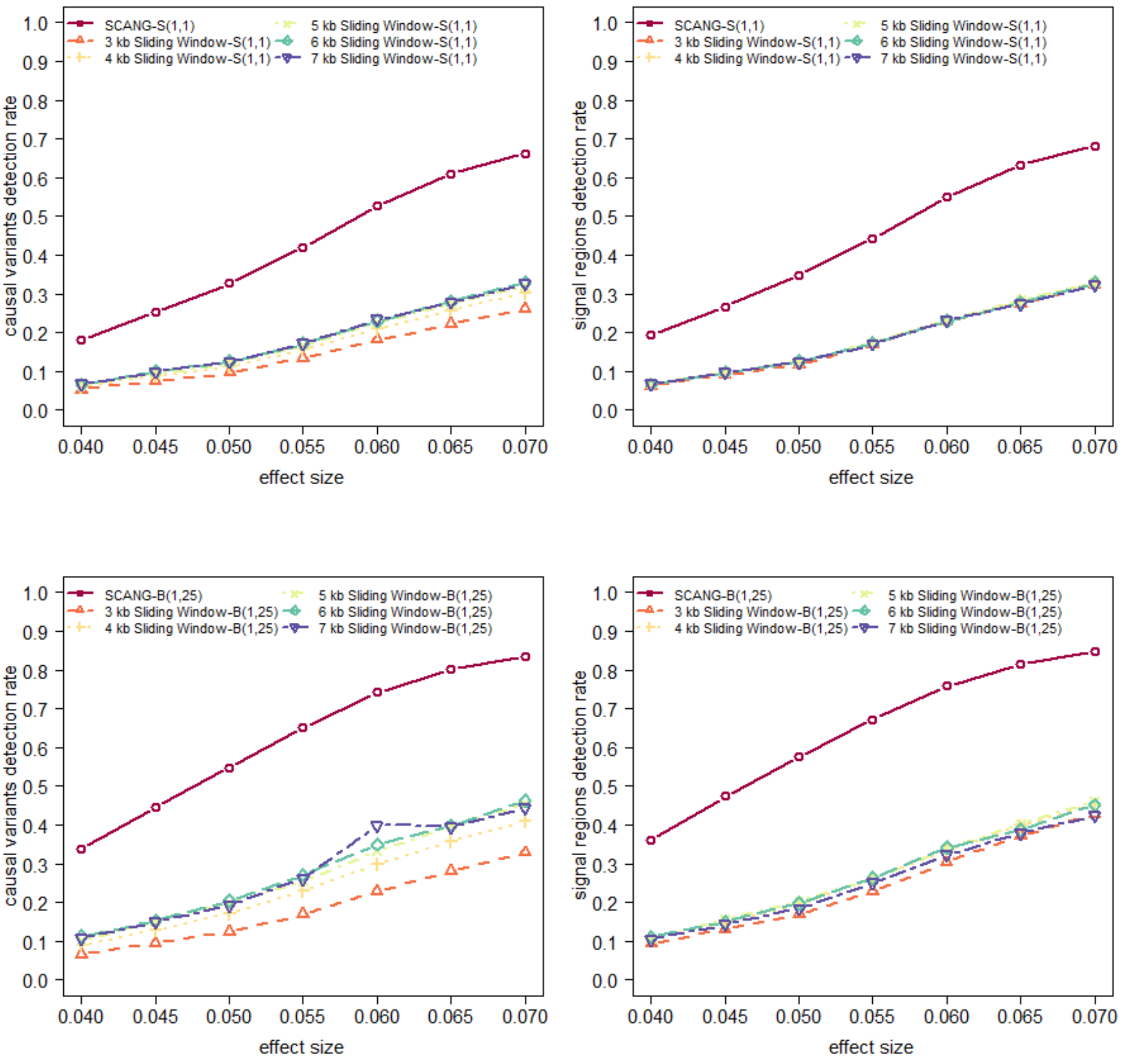

SCANG-S(1,1)
3 kb Sliding Window-S(1,1)
4 kb Sliding Window-S(1,1)
5 kb Sliding Window-S(1,1)
6 kb Sliding Window-S(1,1)
7 kb Sliding Window-S(1,1)
causal variants detection rate
signal regions detection rate
effect size
SCANG-B(1,25)
3 kb Sliding Window-B(1,25)
4 kb Sliding Window-B(1,25)
5 kb Sliding Window-B(1,25)
6 kb Sliding Window-B(1,25)
7 kb Sliding Window-B(1,25)
0.0 0.1 0.2 0.3 0.4 0.5 0.6 0.7 0.8 0.9 1.0
0.040 0.045 0.050 0.055 0.060 0.065 0.070

**Figure S13. Power comparisons of SCANG and the corresponding sliding window procedures using SKAT(1,25) and Burden(1,1) for binary trait analysis with multiple effect sizes when 10% of rare variants are causal variants**

Empirical power was evaluated by causal variants detection rate and signal region detection rate defined in the simulation section. Both two criteria were calculated at the genome-wise/family-wise type I error $\alpha = 0.05$. We simulated **10%** of the rare variants within random 3 kb-6 kb regions to be causal. We set for **binary phenotypes** with the effect size $\beta_j = c|\,log_{10} MAF_j\,|$ where the effect sizes decrease with MAFs, and the maximum log odds ratio with $\mathrm{MAF} = 5 \times 10^{-5}$ varies from $\log(2)$ to $\log(4.4)$. The coefficients for the causal variants were all positive. The causal variants detection rate (left panel) is the proportion of detected causal variants in 2,000 simulated whole genome data sets, where a causal variant is called detected if it is in one of the detected signal regions. The signal region detection rate (right panel) is the proportion of detected signal regions in 2,000 simulated whole genome data sets, where a signal region is called detected if it is overlapped with one of the detected signal regions. For each configuration, the total sample size considered were 10,000. For each setting, six methods were compared: SCANG and sliding window procedures using **SKAT(1,25)** and **Burden(1,1)** with searching window length used in the sliding window method equal to 3 kb, 4 kb, 5 kb, 6 kb and 7 kb. SKAT(1,25) and Burden(1,1) are denoted by S(1,25) and B(1,1), and the corresponding SCANG methods are denoted by SCANG-S(1,25) and SCANG-B(1,1) (the two numbers in the parentheses are the values of beta(MAF) weight parameters $a_1$ and $a_2$, respectively). For SCANG, the range of search window lengths was set by the numbers of variants in searching windows between the 1%st percentile of the numbers of variants of all 3 kb sliding windows and the 99%th percentile of the numbers of variants of all 7 kb sliding windows.

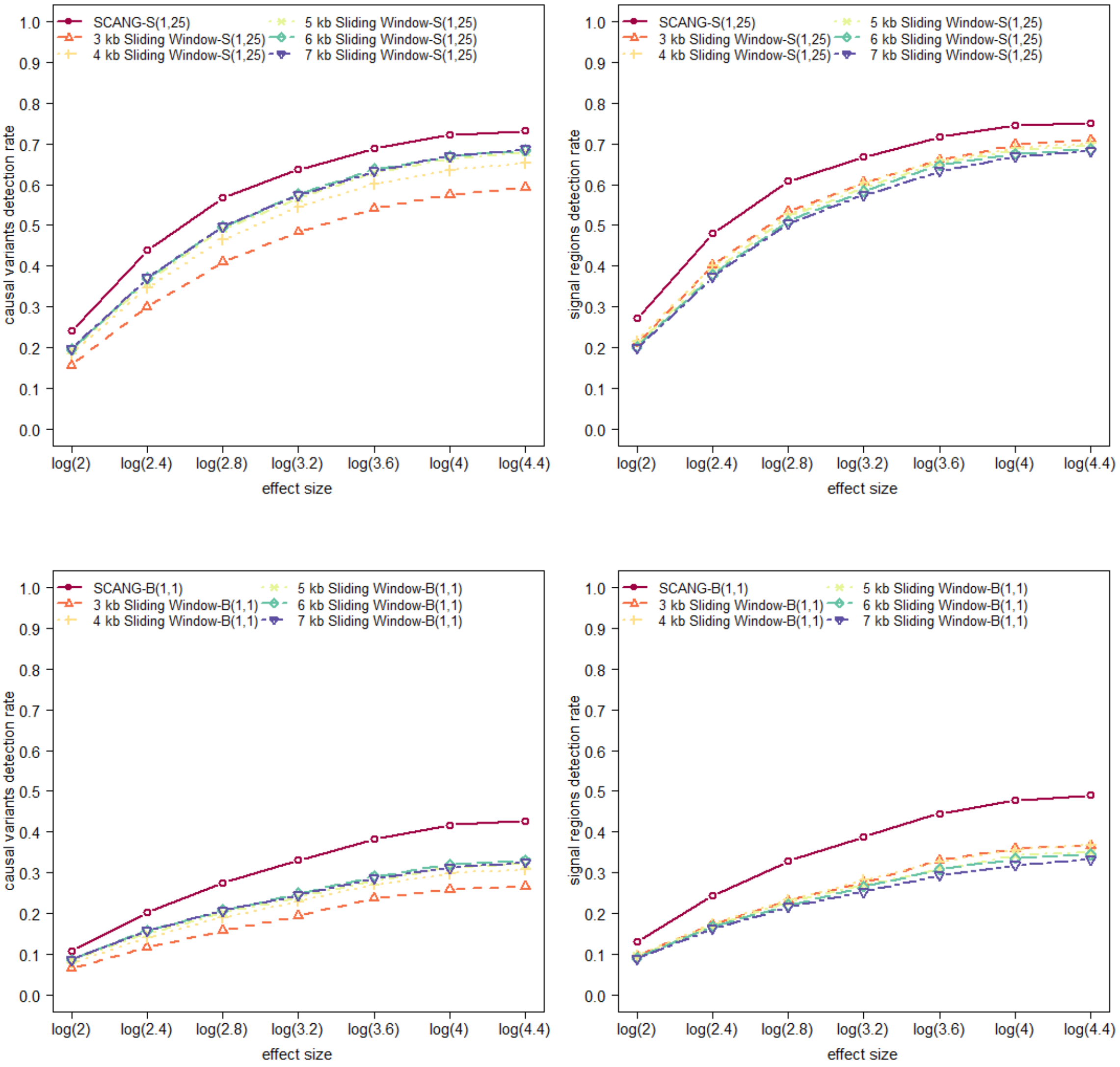
SCANG-S(1,25)
3 kb Sliding Window-S(1,25)
4 kb Sliding Window-S(1,25)
5 kb Sliding Window-S(1,25)
6 kb Sliding Window-S(1,25)
7 kb Sliding Window-S(1,25)
causal variants detection rate
signal regions detection rate
SCANG-B(1,1)
3 kb Sliding Window-B(1,1)
4 kb Sliding Window-B(1,1)
5 kb Sliding Window-B(1,1)
6 kb Sliding Window-B(1,1)
7 kb Sliding Window-B(1,1)
0.0
0.1
0.2
0.3
0.4
0.5
0.6
0.7
0.8
0.9
1.0
log(2)
log(2.4)
log(2.8)
log(3.2)
log(3.6)
log(4)
log(4.4)
effect size

**Figure S14**. **Power comparisons of SCANG and the corresponding sliding window procedures using SKAT(1,1) and Burden(1,25) for binary trait analysis with multiple effect sizes when 10% of rare variants are causal variants**

Empirical power was evaluated by causal variants detection rate and signal region detection rate defined in the simulation section. Both two criteria were calculated at the genome-wise/family-wise type I error $\alpha = 0.05$. We simulated **10%** of the rare variants within random 3 kb-6 kb regions to be causal. We set for **binary phenotypes** with the effect size $\beta_j = c|\, log_{10} MAF_j \,|$ where the effect sizes decrease with MAFs, and the maximum log odds ratio with $\mathrm{MAF} = 5 \times 10^{-5}$ varies from $\log(2)$ to $\log(4.4)$. The coefficients for the causal variants were all positive. The causal variants detection rate (left panel) is the proportion of detected causal variants in 2,000 simulated whole genome data sets, where a causal variant is called detected if it is in one of the detected signal regions. The signal region detection rate (right panel) is the proportion of detected signal regions in 2,000 simulated whole genome data sets, where a signal region is called detected if it is overlapped with one of the detected signal regions. For each configuration, the total sample size considered were 10,000. For each setting, six methods were compared: SCANG and sliding window procedures using **SKAT(1,1)** and **Burden(1,25)** with searching window length used in the sliding window method equal to 3 kb, 4 kb, 5 kb, 6 kb and 7 kb. SKAT(1,1) and Burden(1,25) are denoted by S(1,1) and B(1,25), and the corresponding SCANG methods are denoted by SCANG-S(1,1) and SCANG-B(1,25) (the two numbers in the parentheses are the values of beta(MAF) weight parameters $a_1$ and $a_2$, respectively). For SCANG, the range of search window lengths was set by the numbers of variants in searching windows between the 1%st percentile of the numbers of variants of all 3 kb sliding windows and the 99%th percentile of the numbers of variants of all 7 kb sliding windows.

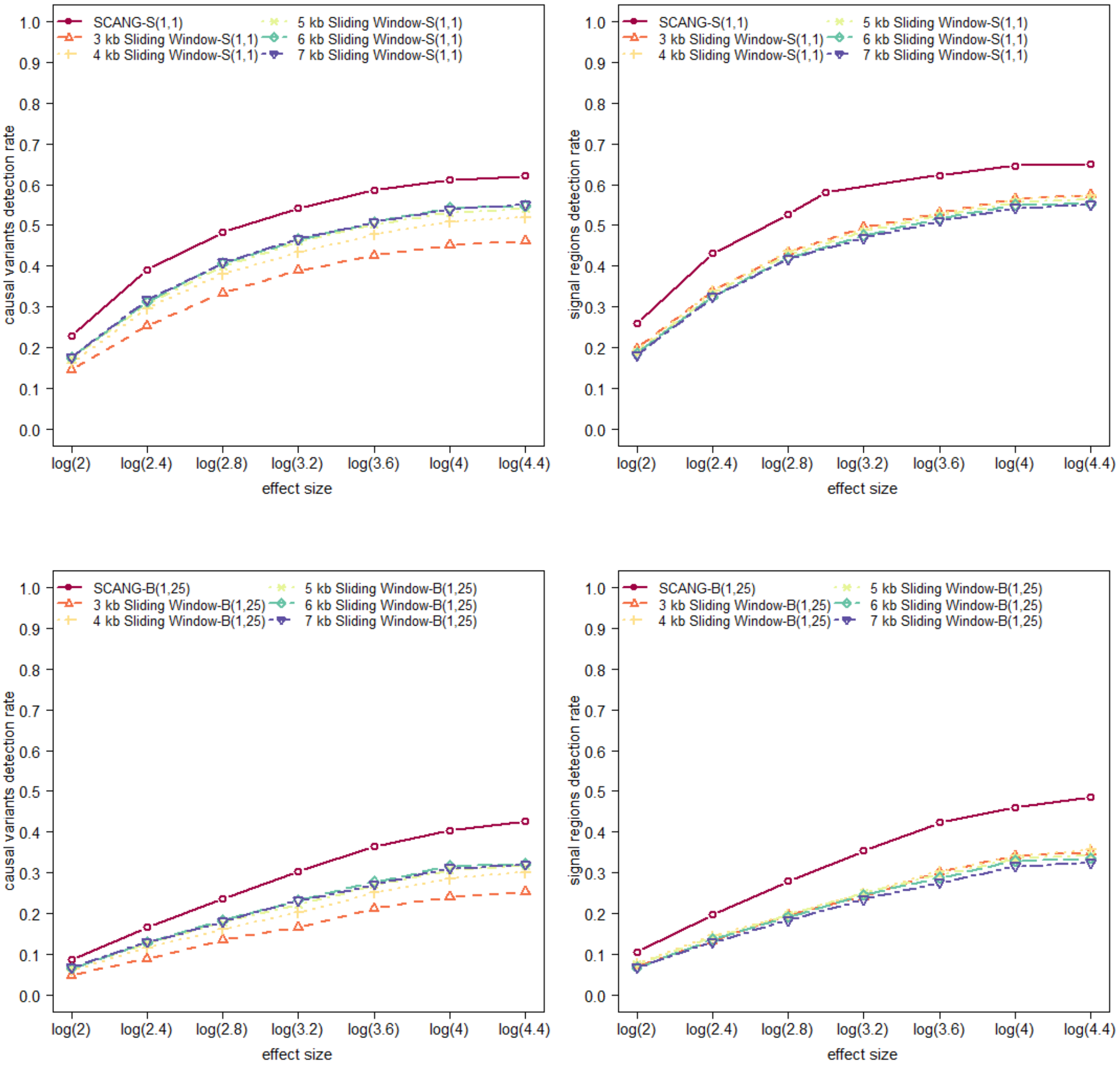
SCANG-S(1,1)
3 kb Sliding Window-S(1,1)
4 kb Sliding Window-S(1,1)
5 kb Sliding Window-S(1,1)
6 kb Sliding Window-S(1,1)
7 kb Sliding Window-S(1,1)
causal variants detection rate
signal regions detection rate
effect size
log(2)
log(2.4)
log(2.8)
log(3.2)
log(3.6)
log(4)
log(4.4)
SCANG-B(1,25)
3 kb Sliding Window-B(1,25)
4 kb Sliding Window-B(1,25)
5 kb Sliding Window-B(1,25)
6 kb Sliding Window-B(1,25)
7 kb Sliding Window-B(1,25)

**Figure S15**. **Power comparisons of SCANG and the corresponding sliding window procedures using SKAT(1,25) and Burden(1,1) for binary trait analysis with multiple effect sizes when 40% of rare variants are causal variants**

Empirical power was evaluated by causal variants detection rate and signal region detection rate defined in the simulation section. Both two criteria were calculated at the genome-wise/family-wise type I error $\alpha = 0.05$. We simulated **40%** of the rare variants within random 3 kb-6 kb regions to be causal. We set for **binary phenotypes** with the effect size $\beta_j = c|\,log_{10} MAF_j\,|$ where the effect sizes decrease with MAFs, and the maximum log odds ratio with $\mathrm{MAF} = 5 \times 10^{-5}$ varies from $\log(1.4)$ to $\log(1.7)$. The coefficients for the causal variants were all positive. The causal variants detection rate (left panel) is the proportion of detected causal variants in 2,000 simulated whole genome data sets, where a causal variant is called detected if it is in one of the detected signal regions. The signal region detection rate (right panel) is the proportion of detected signal regions in 2,000 simulated whole genome data sets, where a signal region is called detected if it is overlapped with one of the detected signal regions. For each configuration, the total sample size considered were 10,000. For each setting, six methods were compared: SCANG and sliding window procedures using **SKAT(1,25)** and **Burden(1,1)** with searching window length used in the sliding window method equal to 3 kb, 4 kb, 5 kb, 6 kb and 7 kb. SKAT(1,25) and Burden(1,1) are denoted by S(1,25) and B(1,1), and the corresponding SCANG methods are denoted by SCANG-S(1,25) and SCANG-B(1,1) (the two numbers in the parentheses are the values of beta(MAF) weight parameters $a_1$ and $a_2$, respectively). For SCANG, the range of search window lengths was set by the numbers of variants in searching windows between the 1%st percentile of the numbers of variants of all 3 kb sliding windows and the 99%th percentile of the numbers of variants of all 7 kb sliding windows.

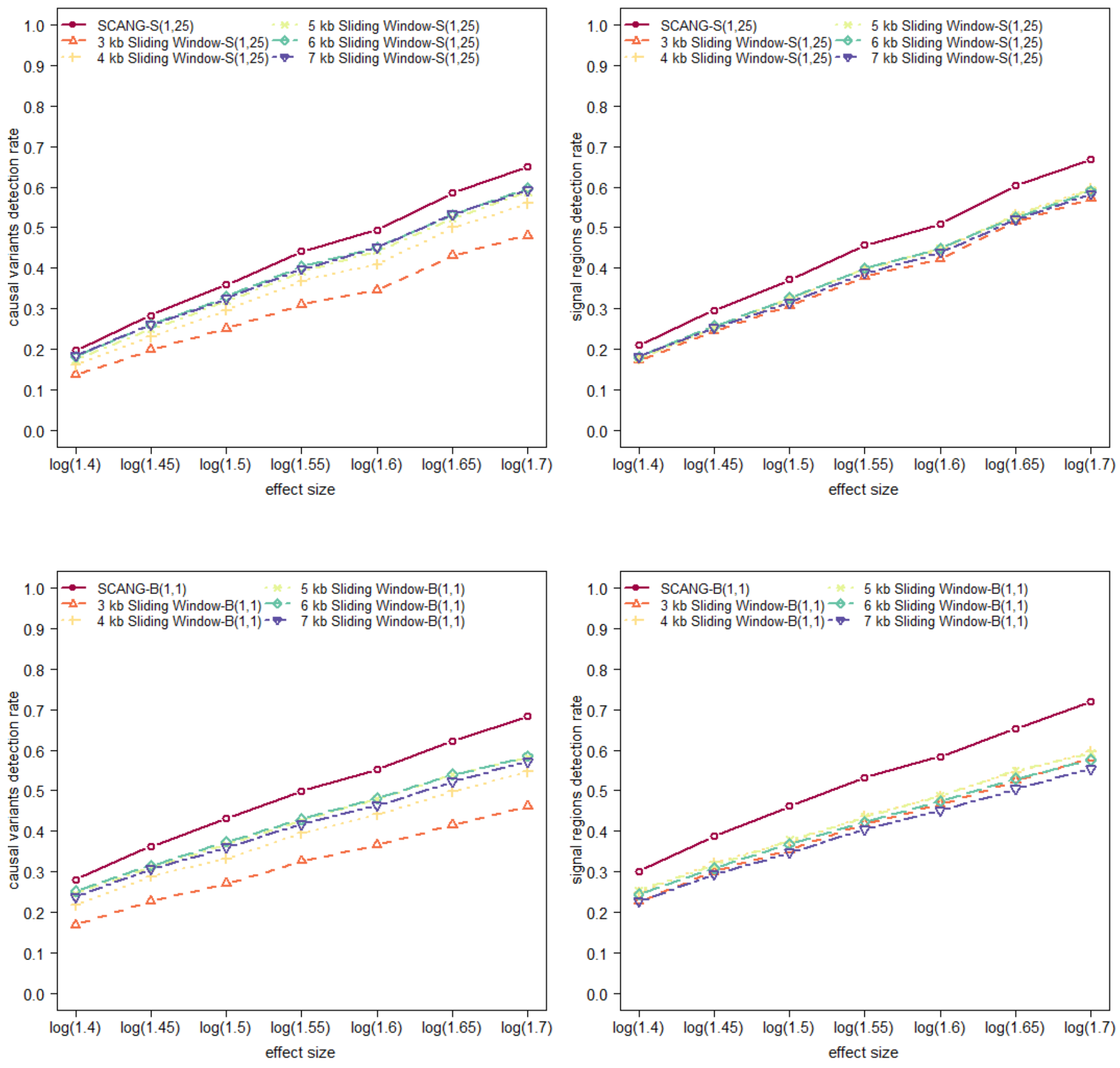

SCANG-S(1,25)
3 kb Sliding Window-S(1,25)
4 kb Sliding Window-S(1,25)
5 kb Sliding Window-S(1,25)
6 kb Sliding Window-S(1,25)
7 kb Sliding Window-S(1,25)
causal variants detection rate
signal regions detection rate
effect size
log(1.4)
log(1.45)
log(1.5)
log(1.55)
log(1.6)
log(1.65)
log(1.7)
SCANG-B(1,1)
3 kb Sliding Window-B(1,1)
4 kb Sliding Window-B(1,1)
5 kb Sliding Window-B(1,1)
6 kb Sliding Window-B(1,1)
7 kb Sliding Window-B(1,1)

**Figure S16**. . **Power comparisons of SCANG and the corresponding sliding window procedures using SKAT(1,1) and Burden(1,25) for binary trait analysis with multiple effect sizes when 40% of rare variants are causal variants**

Empirical power was evaluated by causal variants detection rate and signal region detection rate defined in the simulation section. Both two criteria were calculated at the genome-wise/family-wise type I error $\alpha = 0.05$. We simulated **40%** of the rare variants within random 3 kb-6 kb regions to be causal. We set for **binary phenotypes** with the effect size $\beta_j = c|\, log_{10} MAF_j\, |$ where the effect sizes decrease with MAFs, and the maximum log odds ratio with $\mathrm{MAF} = 5 \times 10^{-5}$ varies from $\log(1.4)$ to $\log(1.7)$. The coefficients for the causal variants were all positive. The causal variants detection rate (left panel) is the proportion of detected causal variants in 2,000 simulated whole genome data sets, where a causal variant is called detected if it is in one of the detected signal regions. The signal region detection rate (right panel) is the proportion of detected signal regions in 2,000 simulated whole genome data sets, where a signal region is called detected if it is overlapped with one of the detected signal regions. For each configuration, the total sample size considered were 10,000. For each setting, six methods were compared: SCANG and sliding window procedures using **SKAT(1,1)** and **Burden(1,25)** with searching window length used in the sliding window method equal to 3 kb, 4 kb, 5 kb, 6 kb and 7 kb. SKAT(1,1) and Burden(1,25) are denoted by S(1,1) and B(1,25), and the corresponding SCANG methods are denoted by SCANG-S(1,1) and SCANG-B(1,25) (the two numbers in the parentheses are the values of beta(MAF) weight parameters $a_1$ and $a_2$, respectively). For SCANG, the range of search window lengths was set by the numbers of variants in searching windows between the 1%st percentile of the numbers of variants of all 3 kb sliding windows and the 99%th percentile of the numbers of variants of all 7 kb sliding windows.

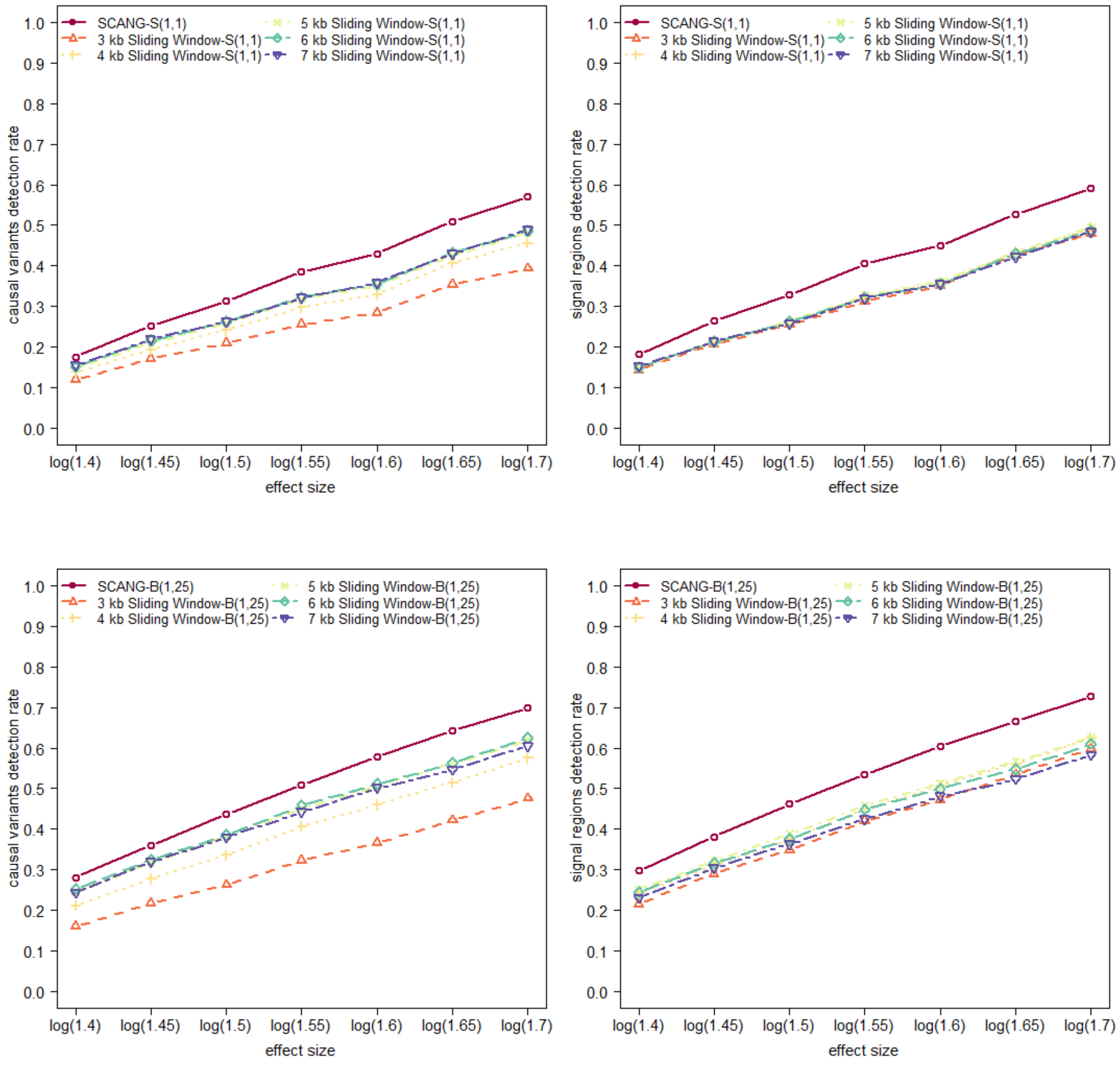

SCANG-S(1,1)
3 kb Sliding Window-S(1,1)
4 kb Sliding Window-S(1,1)
5 kb Sliding Window-S(1,1)
6 kb Sliding Window-S(1,1)
7 kb Sliding Window-S(1,1)
causal variants detection rate
signal regions detection rate
effect size
log(1.4)
log(1.45)
log(1.5)
log(1.55)
log(1.6)
log(1.65)
log(1.7)
0.0
0.1
0.2
0.3
0.4
0.5
0.6
0.7
0.8
0.9
1.0
SCANG-B(1,25)
3 kb Sliding Window-B(1,25)
4 kb Sliding Window-B(1,25)
5 kb Sliding Window-B(1,25)
6 kb Sliding Window-B(1,25)
7 kb Sliding Window-B(1,25)

**Figure S17**. **Power comparison of SCANG-O, SCANG-S(1,1), SCANG-S(1,25), SCANG-B(1,1) and SCANG-B(1,25) for binary trait analysis**

SKAT($a_1, a_2$) and Burden($a_1, a_2$) are denoted by S($a_1, a_2$) and B($a_1, a_2$), and the corresponding SCANG methods are denoted by SCANG-S($a_1, a_2$) and SCANG-B($a_1, a_2$). The two numbers in the parentheses are the values of beta(MAF) weight parameters $a_1$ and $a_2$, respectively. Empirical power was evaluated by causal variants detection rate and signal region detection rate and both two criteria were calculated at the genome-wise/family-wise type I error $\alpha = 0.05$. We simulated 10% and 40% of the rare variants within random 3 kb-6 kb regions to be causal. We set for binary phenotypes with the maximum odds ratio equal to 3 and 1.65 for variants with $\mathrm{MAF} = 5 \times 10^{-5}$ when 10% and 40% causal variants, respectively, and let the effect sizes decrease with MAFs $\beta_j = c|\,log_{10} MAF_j\,|$. The coefficients for the causal variants were all positive. The causal variants detection rate is the proportion of detected causal variants in 2,000 simulated data sets, where a causal variant is called detected if it is in one of the detected signal regions. The signal region detection rate is the proportion of detected signal regions in 2,000 simulated data sets, where a signal region is called detected if it is overlapped with one of the detected signal regions. The range of search window lengths was set by the numbers of variants in searching windows between the 1%st percentile of the numbers of variants of all 3 kb sliding windows and the 99%th percentile of the numbers of variants of all 7 kb sliding windows. For each configuration, the total sample size considered were 2,500, 5,000 and 10,000. (A) Causal variants detection rate when 10% causal variants. (B) Causal variants detection rate when 40% causal variants. (C) Signal region detection rate when 10% causal variants. (D) Signal region detection rate when 40% causal variants.

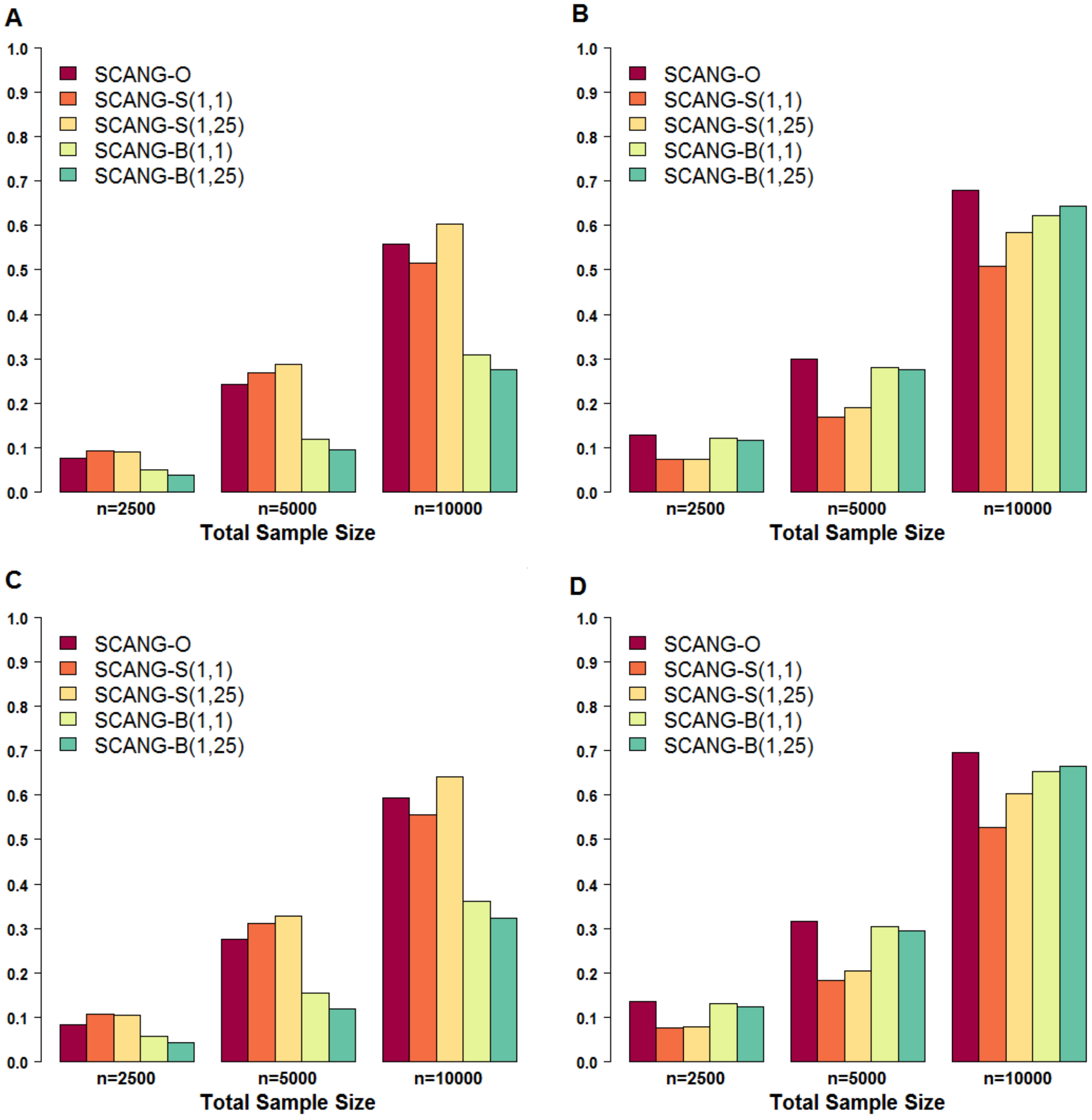

A
B
C
D
1.0
0.9
0.8
0.7
0.6
0.5
0.4
0.3
0.2
0.1
0.0
SCANG-O
SCANG-S(1,1)
SCANG-S(1,25)
SCANG-B(1,1)
SCANG-B(1,25)
n=2500
n=5000
n=10000
Total Sample Size

**Figure S18**. **Power comparison of SCANG-O, SCANG-S(1,1), SCANG-S(1,25), SCANG-B(1,1) and SCANG-B(1,25) for continuous trait analysis with multiple effect sizes**

Empirical power from simulation studies evaluated using causal variant detection rate and signal region detection rate using SCANG-O, SCANG-S(1,1), SCANG-S(1,25), SCANG-B(1,1) and SCANG-B(1,25) (the two numbers in the parentheses are the values of beta(MAF) weight parameters $a_1$ and $a_2$, respectively in **continuous trait** analysis for **multiple effect sizes** of 10,000 samples. Empirical power was calculated at the genome-wise/family-wise type I error $\alpha = 0.05$. We simulated 10% and 40% of the rare variants within random 3 kb-6 kb regions to be causal. We set the effect sizes decrease with MAFs and let $\beta_j = c| \, log_{10} MAF_j \, |$. For 10% causal variants, $c$ varies from 0.11 to 0.23, and for 40% causal variants, $c$ varies from 0.04 to 0.07. The causal variants detection rate is the proportion of detected causal variants in 2,000 simulated whole genome data sets, where a causal variant is called detected if it is in one of the detected signal regions. The signal region detection rate is the proportion of detected signal regions in 2,000 simulated whole genome data sets, where a signal region is called detected if it is overlapped with one of the detected signal regions. The range of search window lengths was set by the numbers of variants in searching windows between the 1%st percentile of the numbers of variants of all 3 kb sliding windows and the 99%th percentile of the numbers of variants of all 7 kb sliding windows. (A) Causal variants detection rate when 10% causal variants. (B) Causal variants detection rate when 40% causal variants. (C) Signal regions detection rate when 10% causal variants. (D) Signal regions detection rate when 40% causal variants.

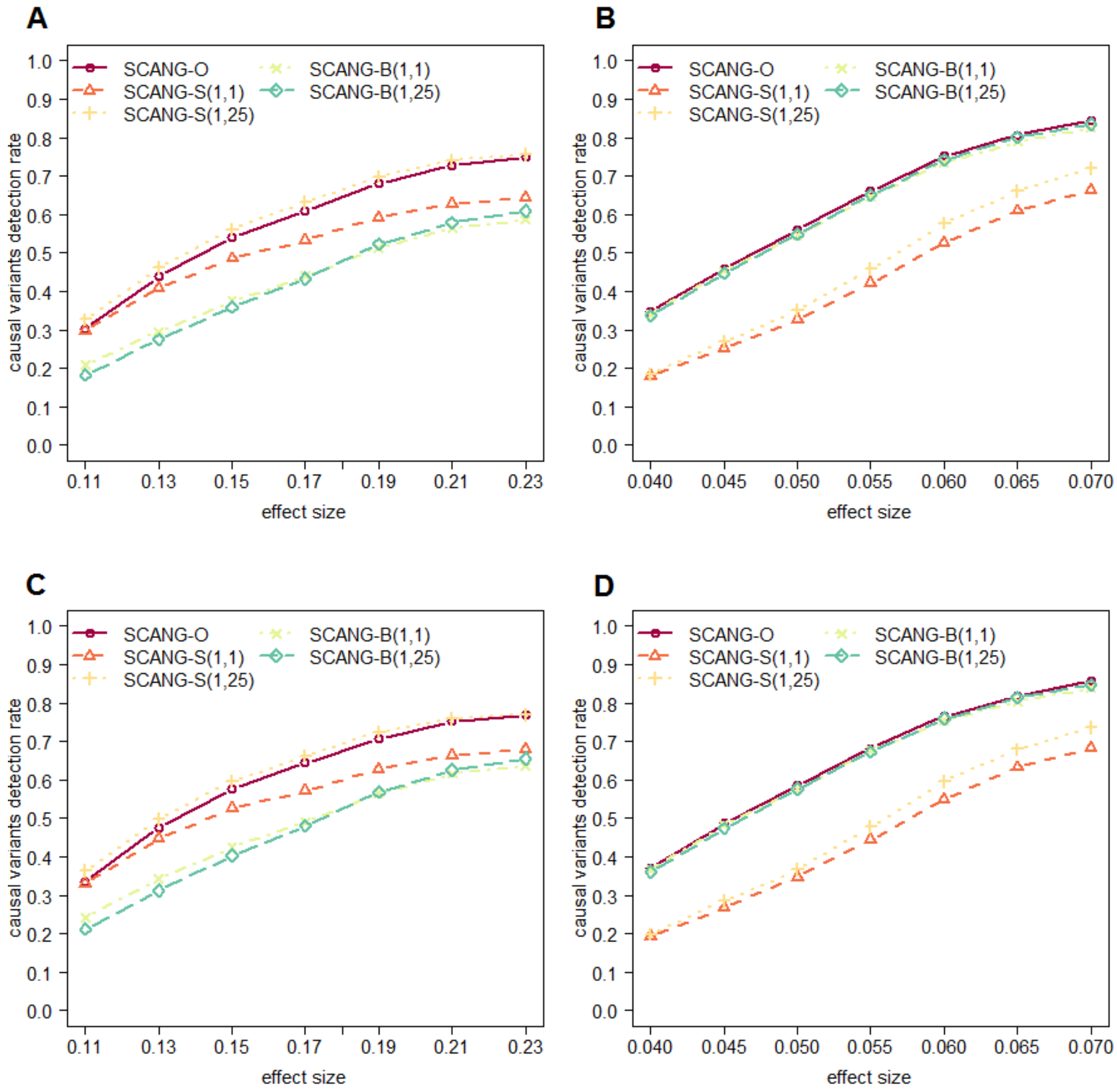
A
B
C
D
SCANG-O
SCANG-S(1,1)
SCANG-S(1,25)
SCANG-B(1,1)
SCANG-B(1,25)
causal variants detection rate
effect size
0.0
0.1
0.2
0.3
0.4
0.5
0.6
0.7
0.8
0.9
1.0
0.11
0.13
0.15
0.17
0.19
0.21
0.23
0.040
0.045
0.050
0.055
0.060
0.065
0.070

**Figure S19**. **Power comparison of SCANG-O, SCANG-S(1,1), SCANG-S(1,25), SCANG-B(1,1) and SCANG-B(1,25) for binary trait analysis with multiple effect sizes** Empirical power from simulation studies evaluated using causal variant detection rate and signal region detection rate using SCANG-O, SCANG-S(1,1), SCANG-S(1,25), SCANG-B(1,1) and SCANG-B(1,25) (the two numbers in the parentheses are the values of beta(MAF) weight parameters $a_1$ and $a_2$, respectively in **binary trait** analysis for **multiple effect sizes** of 10,000 samples. Empirical power was calculated at the genome-wise/family-wise type I error $\alpha = 0.05$. We simulated 10% and 40% of the rare variants within random 3 kb-6 kb regions to be causal. We set the effect sizes decrease with MAFs and let $\beta_j = c| log_{10} MAF_j |$. For 10% causal variants, the maximum log odds ratio with $\mathrm{MAF} = 5 \times 10^{-5}$ varies from $\log(2)$ to $\log(4.4)$, and for 40% causal variants, the maximum log odds ratio with $\mathrm{MAF} = 5 \times 10^{-5}$ varies from $\log(1.4)$ to $\log(1.7)$. The causal variants detection rate is the proportion of detected causal variants in 2,000 simulated whole genome data sets, where a causal variant is called detected if it is in one of the detected signal regions. The signal region detection rate is the proportion of detected signal regions in 2,000 simulated whole genome data sets, where a signal region is called detected if it is overlapped with one of the detected signal regions. The range of search window lengths was set by the numbers of variants in searching windows between the 1%st percentile of the numbers of variants of all 3 kb sliding windows and the 99%th percentile of the numbers of variants of all 7 kb sliding windows. (A) Causal variants detection rate when 10% causal variants. (B) Causal variants detection rate when 40% causal variants. (C) Signal regions detection rate when 10% causal variants. (D) Signal regions detection rate when 40% causal variants.

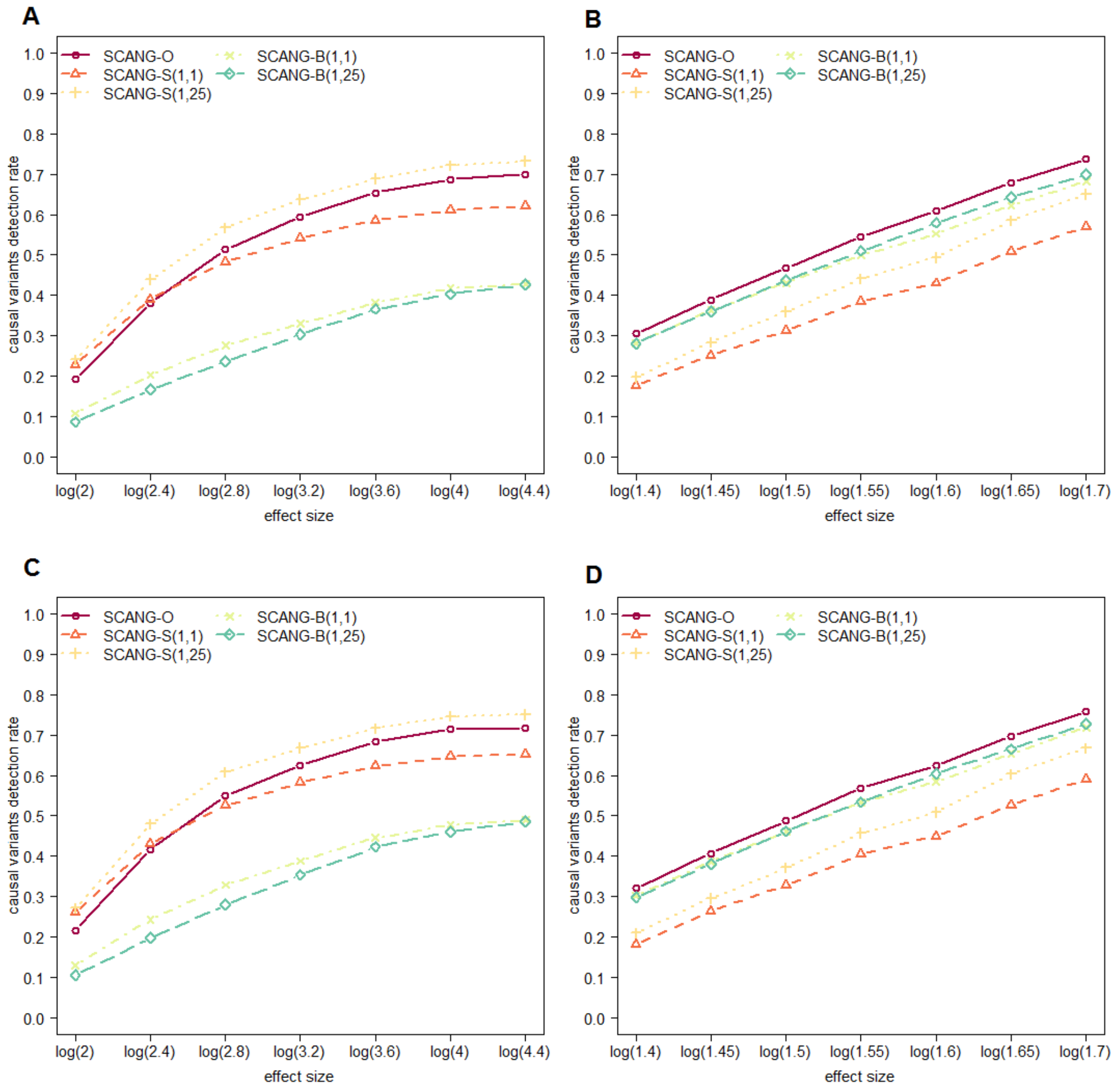

A
B
C
D
SCANG-O
SCANG-S(1,1)
SCANG-S(1,25)
SCANG-B(1,1)
SCANG-B(1,25)
causal variants detection rate
effect size
log(2)
log(2.4)
log(2.8)
log(3.2)
log(3.6)
log(4)
log(4.4)
log(1.4)
log(1.45)
log(1.5)
log(1.55)
log(1.6)
log(1.65)
log(1.7)

**Figure S20**. **Genetic landscape of the windows significantly associated with Lp(a) levels on chromosome 6q25.3-6q26 among European Americans in the ARIC Whole Genome Sequencing Study (n=1,705)**

Five methods are compared: SCANG and 4 kb sliding window procedures using SKAT(1,25) and Burden(1,1) (denoted by S(1,25) and B(1,1) respectively), with the corresponding SCANG methods denoted by SCANG-S(1,25) and SCANG-B(1,1) (the two numbers in the parentheses are the values of beta(MAF) weight parameters $a_1$ and $a_2$, respectively), and SCANG-O, which used the proposed omnibus test that aggregated SKAT(1,1), SKAT(1,25), Burden(1,1) and Burden(1,25) using ACAT in the SCANG framework. A dot means that the sliding window at this location is significant using the method that the color of the dot represents.

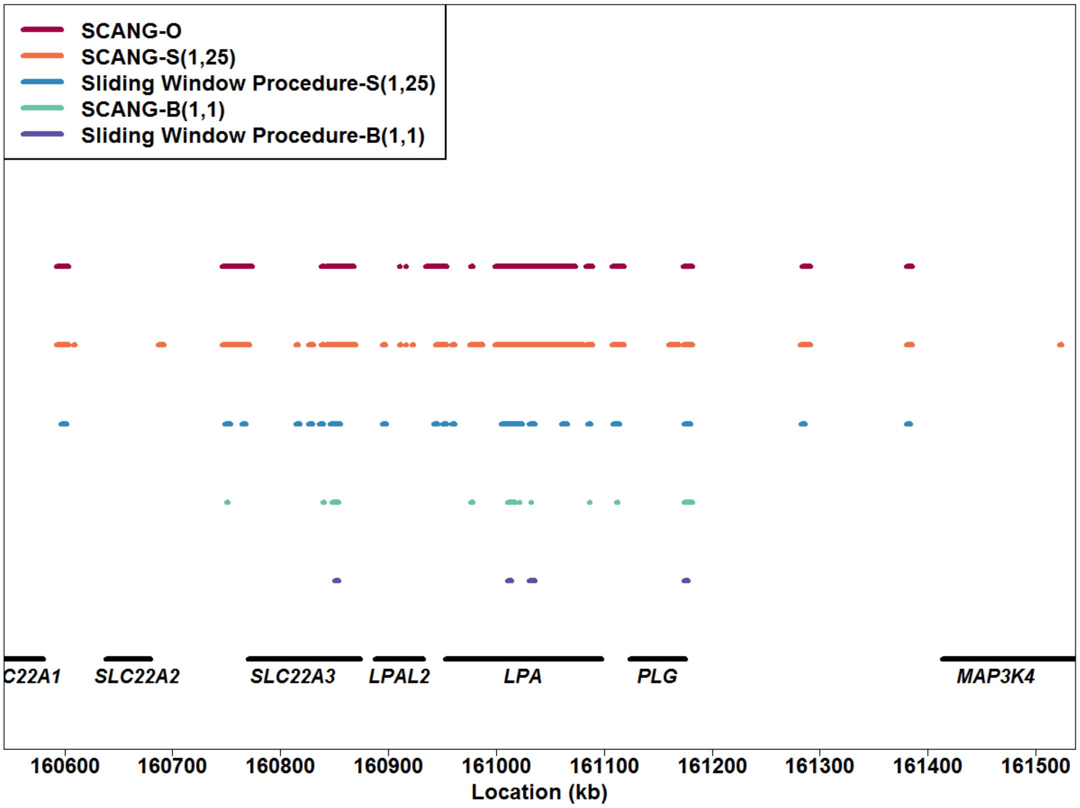

**Figure S21. Toy example of searching algorithm for multiple signal regions**

Step 1. Calculate the set-based p-value $p(I)$ for intervals with number of variants satisfies $L_{min} \leq |I| \leq L_{max}$.

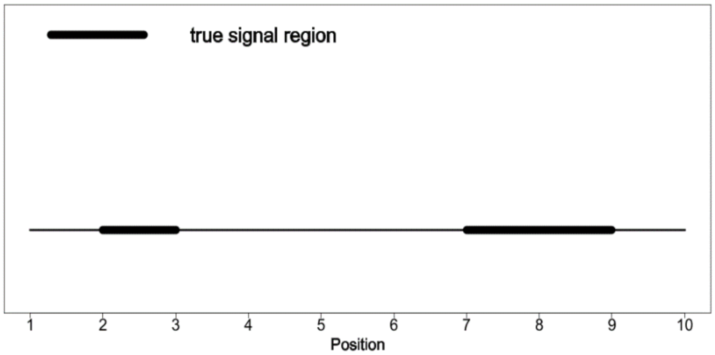


Step 2. Find candidate regions with p-value $(I) < h(\alpha, L_{min}, L_{max})$ . Select the candidate region with smallest $p(I)$ as estimated signal region.

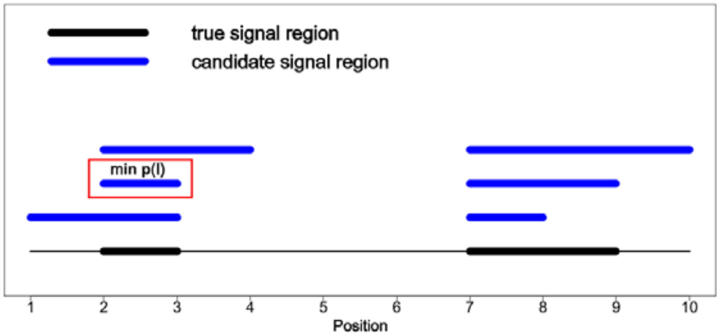


Step 3. Remove the candidate regions which overlaps by more than $f$ with the estimated region. Then again select the candidate region with smallest $p(I)$ as the next estimated signal region.

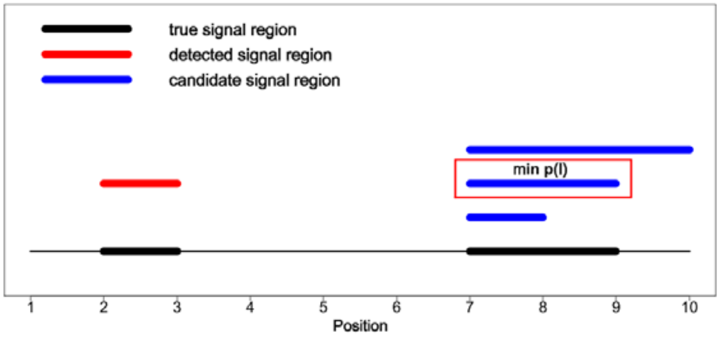


Step 4. Repeat Step 3 until there is no candidate region.

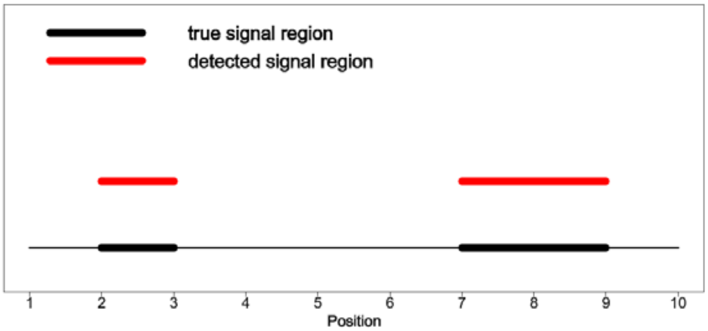

**Table S1**. **Genome-wise/family-wise empirical type I error rates from simulation studies using SCANG-B(1,1), SCANG-B(1,25), SCANG-S(1,1) , SCANG-S(1,25) and SCANG-O at the genome-wide/family-wise significance level of α=0.05 and 0.01 when $L_{min} = 20$ and $L_{max} = 120$**

SKAT(1,25) and Burden(1,1) are denoted by S(1,25) and B(1,1), and the corresponding SCANG methods are denoted by SCANG-S(1,25) and SCANG-B(1,1) (the two numbers in the parentheses are the values of beta(MAF) weight parameters $a_1$ and $a_2$, respectively. The total sample size *n* was 2,500, 5,000 and 10,000. The smallest number of variants in searching windows are $L_{min} = 20$ and the corresponding largest numbers of variants in searching windows are $L_{max} = 120$. Each cell represents the empirical type I error rate estimate that was calculated as the proportion of p-values less than α under the null hypothesis on the basis of $10^4$ replicates.

| | | Continuous Traits | | | Binary Traits | | |
|---|---|---|---|---|---|---|---|
| Total Sample Size | Size | n=2500 | n=5000 | n=10000 | n=2500 | n=5000 | n=10000 |
| SCANG-B(1,1) | 0.05 | 0.0468 | 0.0506 | 0.0459 | 0.0485 | 0.0498 | 0.0480 |
| | 0.01 | 0.0095 | 0.0099 | 0.0087 | 0.0086 | 0.0104 | 0.0094 |
| SCANG-B(1,25) | 0.05 | 0.0463 | 0.0500 | 0.0455 | 0.0461 | 0.0482 | 0.0486 |
| | 0.01 | 0.0081 | 0.0096 | 0.0099 | 0.0095 | 0.0088 | 0.0104 |
| SCANG-S(1,1) | 0.05 | 0.0481 | 0.0498 | 0.0459 | 0.0396 | 0.0435 | 0.0429 |
| | 0.01 | 0.0093 | 0.0093 | 0.0094 | 0.0087 | 0.0090 | 0.0093 |
| SCANG-S(1,25) | 0.05 | 0.0466 | 0.0466 | 0.0458 | 0.0347 | 0.0421 | 0.0412 |
| | 0.01 | 0.0089 | 0.0096 | 0.0093 | 0.0059 | 0.0083 | 0.0093 |
| SCANG-O | 0.05 | 0.0450 | 0.0480 | 0.0468 | 0.0445 | 0.0483 | 0.0476 |
| | 0.01 | 0.0092 | 0.0104 | 0.0099 | 0.0099 | 0.0096 | 0.0100 |

**Table S2**. **Genome-wise/family-wise empirical type I error rates from simulation studies using SCANG-B(1,1), SCANG-B(1,25), SCANG-S(1,1) , SCANG-S(1,25) and SCANG-O at the genome-wide/family-wise significance level of α=0.05 and 0.01 when $(\boldsymbol{L_{min}}, \boldsymbol{L_{max}})$ = (45,135), (50,150) and (70,185) for 2,500, 5,000 and 10,000 individuals**

SKAT(1,25) and Burden(1,1) are denoted by S(1,25) and B(1,1), and the corresponding SCANG methods are denoted by SCANG-S(1,25) and SCANG-B(1,1) (the two numbers in the parentheses are the values of beta(MAF) weight parameters $a_1$ and $a_2$, respectively. The total sample size *n* was 2,500, 5,000 and 10,000. The smallest number of variants in searching windows are $\boldsymbol{L_{min}} =$ **45, 50, 70** for *n* = 2,500, 5,000, 10,000, and the corresponding largest numbers of variants in searching windows are $\boldsymbol{L_{max}} =$ **135, 150, 185**. Each cell represents the empirical type I error rate estimate that was calculated as the proportion of p-values less than α under the null hypothesis on the basis of $10^4$ replicates.

| | | Continuous Traits | | | Binary Traits | | |
|---|---|---|---|---|---|---|---|
| Total Sample Size | Size | n=2500 | n=5000 | n=10000 | n=2500 | n=5000 | n=10000 |
| SCANG-B(1,1) | 0.05 | 0.0478 | 0.0484 | 0.0485 | 0.0480 | 0.0521 | 0.0481 |
| | 0.01 | 0.0092 | 0.0091 | 0.0089 | 0.0098 | 0.0105 | 0.0106 |
| SCANG-B(1,25) | 0.05 | 0.0469 | 0.0473 | 0.0455 | 0.0473 | 0.0463 | 0.0489 |
| | 0.01 | 0.0088 | 0.0091 | 0.0097 | 0.0102 | 0.0097 | 0.0084 |
| SCANG-S(1,1) | 0.05 | 0.0466 | 0.0485 | 0.0468 | 0.0434 | 0.0477 | 0.0466 |
| | 0.01 | 0.0094 | 0.0090 | 0.0088 | 0.0093 | 0.0100 | 0.0085 |
| SCANG-S(1,25) | 0.05 | 0.0462 | 0.0480 | 0.0470 | 0.0376 | 0.0471 | 0.0455 |
| | 0.01 | 0.0075 | 0.0101 | 0.0098 | 0.0068 | 0.0091 | 0.0094 |
| SCANG-O | 0.05 | 0.0465 | 0.0469 | 0.0454 | 0.0447 | 0.0483 | 0.0471 |
| | 0.01 | 0.0088 | 0.0089 | 0.0097 | 0.0088 | 0.0096 | 0.0102 |
